# Unraveling the functional role of the orphan solute carrier, SLC22A24 in the transport of steroid conjugates through metabolomic and genome-wide association studies

**DOI:** 10.1101/648238

**Authors:** Sook Wah Yee, Adrian Stecula, Huan-Chieh Chien, Ling Zou, Elena V. Feofanova, Marjolein van Borselen, Kit Wun Kathy Cheung, Noha A. Yousri, Karsten Suhre, Jason M. Kinchen, Eric Boerwinkle, Roshanak Irannejad, Bing Yu, Kathleen M. Giacomini

**Author notes:** These authors contributed equally to the work (Sook Wah Yee, Adrian Stecula).

## Abstract

Variation in sex hormone levels has wide implications for health and disease. The genes encoding the proteins involved in steroid disposition represent key determinants of interindividual variation in steroid levels and ultimately, their effects. Beginning with metabolomic data from genome-wide association studies (GWAS), we observed that genetic variants in the orphan transporter, SLC22A24 were significantly associated with levels of androsterone glucuronide and etiocholanolone glucuronide (sentinel SNPs p-value <1×10^−30^). In cells over-expressing human or various mammalian orthologs of SLC22A24, we showed that steroid conjugates and bile acids were substrates of the transporter. Phylogenetic, genomic, and transcriptomic analyses suggested that SLC22A24 has a specialized role in the kidney and appears to function in the reabsorption of organic anions, and in particular, anionic steroids. Phenome-wide analysis showed that functional variants of SLC22A24 are associated with human disease such as cardiovascular diseases and acne, which have been linked to dysregulated steroid metabolism. Collectively, these functional genomic studies reveal a previously uncharacterized protein involved in steroid homeostasis, opening up new possibilities for SLC22A24 as a pharmacological target for regulating steroid levels.

**Author Summary:** Steroid hormones, ranging from sex steroids such as testosterone to glucocorticoids play key roles in human health and disease. Accordingly, the identification of the genes and proteins involved in their synthesis, disposition and elimination has been the subject of numerous genetic studies. We have been intrigued by recent studies demonstrating that genetic variants in or near a gene encoding SLC22A24 are strongly associated with steroid levels. SLC22A24 is an orphan transporter with no known ligands and no known biological functions. In this study, we use cellular and computational methods to show that SLC22A24 transports steroid conjugates, bile acids and other dicarboxylic acids. Based on the direction of association of a common stop codon in SLC22A24 with lower levels of steroids, our studies suggest that the transporter functions to reabsorb steroid conjugates in the kidney, a surprising finding, given that conjugation pathways generally function to polar molecules that are readily eliminated by the kidney. The absence of the transporter gene in many species and its presence in higher order primates suggest that SLC22A24 plays a specialized role in steroid homeostasis. Overall, our studies indicate that SLC22A24 functions in the reabsorption of conjugated steroids in the kidney.

## Introduction

Steroid hormones represent major signaling molecules and have been the subject of numerous investigations[1]. Precise regulation of steroid levels is important for human health and disease, which has stimulated many studies focused on the identification and characterization of the genes, proteins and pathways involved in steroid metabolism and disposition. In addition, genome-wide association studies (GWAS) have been performed to identify the polymorphisms in genes that associate with interindividual variation in the levels of steroids, and to discover additional genes that are involved in steroid disposition. In general, multiple GWAS have led to the identification of genes already known to be involved in steroid metabolism, binding and regulation[2–7]. However, one locus, SLC22A24, has puzzled researchers because it has no known role in steroid homeostasis. SLC22A24 encodes an orphan solute carrier (SLC) transporter which has unknown substrates and ligands [6, 8, 9].

The human solute carrier (SLC) transporter superfamily includes approximately 400 proteins grouped into 52 families[10–13]. SLC22A24, an orphan transporter belongs to the SLC22 family, which transports organic ions[14]. In the human genome, the SLC22 family includes 22 members and represents the third largest in the SLC transporterome, behind SLC25 and SLC35 families. The first human member of this family was cloned and characterized in 1997[15, 16], and since then, members of the family were shown to play critical pharmacological roles in mediating the hepatic and renal elimination of a variety of endogenous compounds and drugs (e.g., *SLC22A1* encoding organic cation transporter 1 (OCT1), *SLC22A2* encoding OCT2, *SLC22A6* encoding organic anion transporter 1 (OAT1), and *SLC22A8* encoding OAT3)[17, 18]. Because of their important pharmacologic role, many members of the SLC22 family are recognized as important determinants of drug disposition and mediators of drug-drug interactions[17], and are frequently studied during drug development[12]. Ironically, this family still contains a large cluster of orphan transporters, including SLC22A10, SLC22A14, SLC22A15, SLC22A17, SLC22A18, SLC22A23, SLC22A24, SLC22A25, and SLC22A31[19, 20]. In fact, in spite of their increasing recognition as playing key roles in drug response and toxicity and indeed in human physiology and pathophysiology, close to 75 members of the human SLC superfamily remain classified as orphan transporters, i.e., they have no assigned ligands[10, 11, 21]. A number of factors appear to contribute to this paradox including a non-unified nomenclature and technical challenges involving working with membrane proteins[11].

SLCs that have been characterized to date function in the transmembrane flux of solutes, which include inorganic ions such as heavy metals, as well as organic compounds, such as neurotransmitters and amino acids. Therefore, the key step in understanding the physiological role of these proteins is to identify their endogenous substrates.

Over the years, various functional and molecular approaches have been employed to identify the endogenous ligands of SLC transporters. One robust approach for the identification of substrates for transporters has been the use of genome-wide association studies (GWAS), to identify single nucleotide polymorphisms (SNPs) in SLCs that associate with circulating levels of various metabolites. For example, the discovery of GLUT9 (encoded by *SLC2A9*) to be a uric acid transporter stemmed from a GWAS focused on the identification of genetic variants associated with hyperuricemia and gout[22, 23] and the characterization of SLC16A9 as a carnitine efflux transporter was motivated by a GWAS with metabolomics[24]. Also, tetradecanedioic acid and hexadecanedioic acid were discovered to be naturally occurring substrates of OATP1B1 (encoded by *SLCO1B1*) in a GWAS aimed at identifying metabolomic biomarkers for the transporter[25, 26].

In this study, we used a combination of computational and experimental approaches to uncover endogenous ligands and substrates of SLC22A24. These approaches included radiolabeled substrate and inhibition screens, metabolomic profiling of cells over-expressing SLC22A24, and sequence and structure-based analyses. To provide further support for the endogenous role of the human SLC22A24, we profiled its expression levels in various tissues and cell types. Phylogenetic analysis revealed that the transporter has orthologs in primates, and a few other mammalian species had direct orthologs of SLC22A24, suggesting a specialized role of the transporter. Our work also implicates the transporter in the disposition of drugs and drug metabolites with a steroid scaffold. To our knowledge, this is the first study to identify endogenous and pharmacological substrates of SLC22A24. Further studies are warranted to understand the full physiological and pharmacological roles of the transporter in human and primate biology.

## Results

### GWAS reveal strong associations of SLC22A24 polymorphisms with steroid metabolite levels

Three independent GWAS reveal strong associations of SNPs in SLC22A24 with steroid levels. In particular, the metabolites that reached genome-wide significance included progesterone (p<10^−11^)[6], androsterone glucuronide (sentinel SNP p<10^−50^)[8, 9] and etiocholanolone glucuronide (sentinel SNP p<10^−30^)[8]. The locus zoom plot shows the significance pattern of SNPs in and around *SLC22A24* for the association with androsterone glucuronide (Fig 1A) and etiocholanolone glucuronide (Fig 1B). The SNPs in the vicinity of SLC22A24 have the most significant associations with androsterone glucuronide and etiocholanolone glucuronide in GWAS[8, 9]. The top SNPs for progesterone (rs112295236) and the sentinel SNPs for androsterone glucuronide and etiocholanolone glucuronide (rs78176967, rs151042642)[6, 8, 9] are in the region between SLC22A24 and SLC22A8. These SNPs are in strong linkage disequilibrium (LD), r^2^>0.8, to SNPs within the SLC22A24 gene (Fig 1A and 1B). Interestingly, a nonsense variant in SLC22A24 (rs11231341, Tyr501Ter, c.1503T>G), is strongly associated with the levels of androsterone glucuronide and etiocholanolone glucuronide (p<5×10^−8^) (Fig 1A and 1B). Using LDLink[27], we identified the SNPs in linkage with rs78176967 and rs151042642, and found that among the missense, non-sense, or splice donor variants in SLC22A24 (rs11231341, rs7945121, rs1939749), SLC22A25 (rs6591771) and SLC22A10 (rs1790218, rs1201559), the SLC22A24 p.Tyr501Ter (rs11231341) has the strongest correlation to the top SNPs associated with the steroid glucuronides, with r^2^=0.22, D’=1.0 (Fig 1C). In addition to the published data, we used data from the Atherosclerosis Risk in Communities Study (ARIC) to confirm the direction of the association of the nonsense variant, SLC22A24 p.Tyr501Ter with plasma levels of the two metabolites. In particular, this independent cohort consists of 1375 Caucasians and 573 African Americans with available genotype data and steroid metabolite levels. Primary analysis showed that the major G-allele, encoding the nonsense variant SLC22A24 p.Tyr501Ter, is associated with lower plasma levels of the two steroid glucuronide conjugates, androsterone glucuronide and etiocholanolone glucuronide (Fig 1D, Table 1). Further, association analysis in ARIC study also confirmed weaker associations of other missense variants or nonsense variants in SLC22A25 and SLC22A10 (**S2 Table**). We were puzzled by the observation that the sentinel SNPs, rs78176967 and rs151042642, exhibited stronger associations with steroid levels (p-values range from 1×10^−55^ to 9×10^−11^) than the nonsense SNP, rs11231341 (encoding p.Tyr501Ter, p-values range from 0.0014 - 1×10^−13^) (Table 1B). As the sentinel SNPs, rs78176967 and rs151042642, are intergenic variants with no known or predicted cis-regulatory effects, we suspected that they may tag true causal variants that are linked to them in addition to rs11231341. Searching for publicly available eQTL, we noted that the splice donor variant in SLC22A24, rs1939749, is the top SNP for associations with SLC22A24 transcript levels in kidney eQTL databases[28–30] (**S1 Fig**). In particular, rs1939749, resides at the 3’ junction of exon 1 and could affect splicing between exons 1 and 2. If there is incorrect splicing at the 3’ junction of exon 1, i.e., with the minor allele variant, it will result in a premature stop codon (**S1 Fig**). We hypothesized that rs1939749 drives the SLC22A24 expression levels and thus the minor allele resulting in lower transcript levels (**S1 Fig**). Interestingly, the A alleles (minor alleles) of the sentinel SNPs (rs78176967 and rs151042642) were in complete linkage disequilibrium (D’ = 1) with the major (G-allele) of rs1939749 and the T-allele of rs11231341 encoding 501Tyr (Fig 1E). The resultant haplotype, rs78176967:rs151042642:rs11231341:rs1939749, = A:A:T:G, which is present at ∼5% allele frequency is expected to retain function as there is no stop codon present at position 501 (allele-and no defective splicing (allele-G). The weaker p-value of the allele encoding the p.Tyr501, rs11231341, is due to the fact that it may occur in two distinct haplotypes, one that is a loss of function due to the allele that is predicted to disrupt splicing (rs1939749), and one that retains function (no splicing defect) (Fig 1E). Overall, these results led to further studies designed to explore the functional role of SLC22A24 as a transporter of steroids and steroid conjugates.

**Fig 1.**
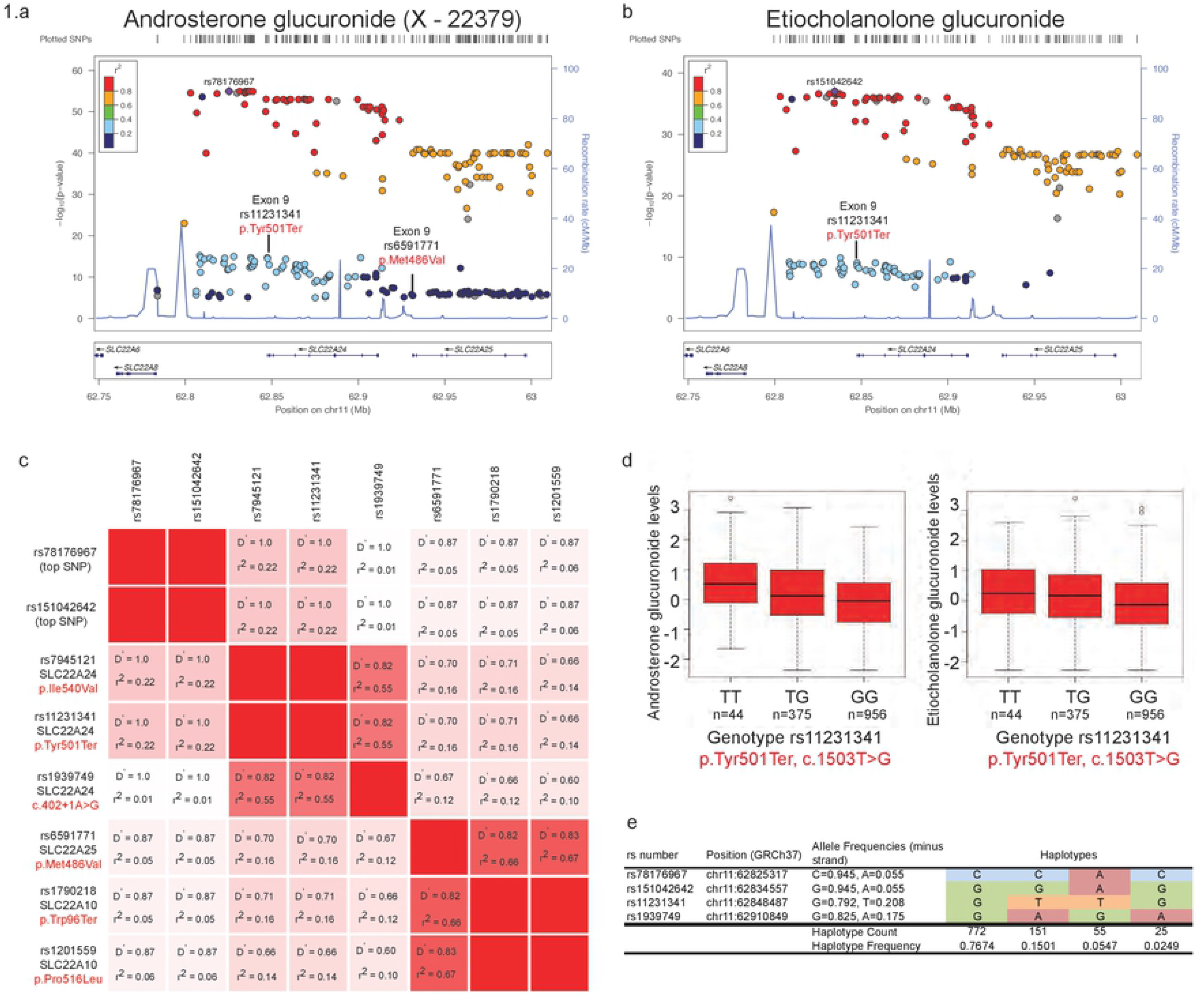
Genetic associations of variants in the *SLC22A24* locus with steroid glucuronide levels. LocusZoom plot showing association results of top variants in the *SLC22A24* locus with **A**) androsterone glucuronide (X-22379) and **B**) etiocholanolone glucuronide, linkage disequilibrium (LD) and recombination rates. The LD estimates are color coded as a heatmap from dark blue (0<r^2^<0.2) to red (0.8<r^2^<1.0). Recombination rate are indicated by the solid blue lines (recombination rate in cM/Mb from HapMap). The bottom panel shows the genes and their orientation. The associations were plotted using available data from Long T. et al. (http://www.hli-opendata.com/Metabolome/, **S1 Table**). **C**) The linkage disequilibrium as calculated by D prime (D’) and correlations (r^2^) between the top SNPs rs781769767 and rs151042642 with missense and nonsense variants in *SLC22A24*, *SLC22A25* and *SLC22A10*. **D**) Genetic association of *SLC22A24* p.Tyr501Ter with normalized serum levels of androsterone glucuronide and etiocholanolone glucuronide in the ARIC study (N=1375 Caucasians). The major allele of rs11231341 coding for p.Tyr501Ter is significantly associated with lower serum levels of androsterone glucuronide and etiocholanolone glucuronide (see Table 1). **E**) Haplotype frequencies of the haplotypes observed for the list of query variants in European population from Phase 3 of the 1000 Genomes Project. This plot is created using LDhap https://ldlink.nci.nih.gov/?tab=ldhap[27]

**Table 1.**
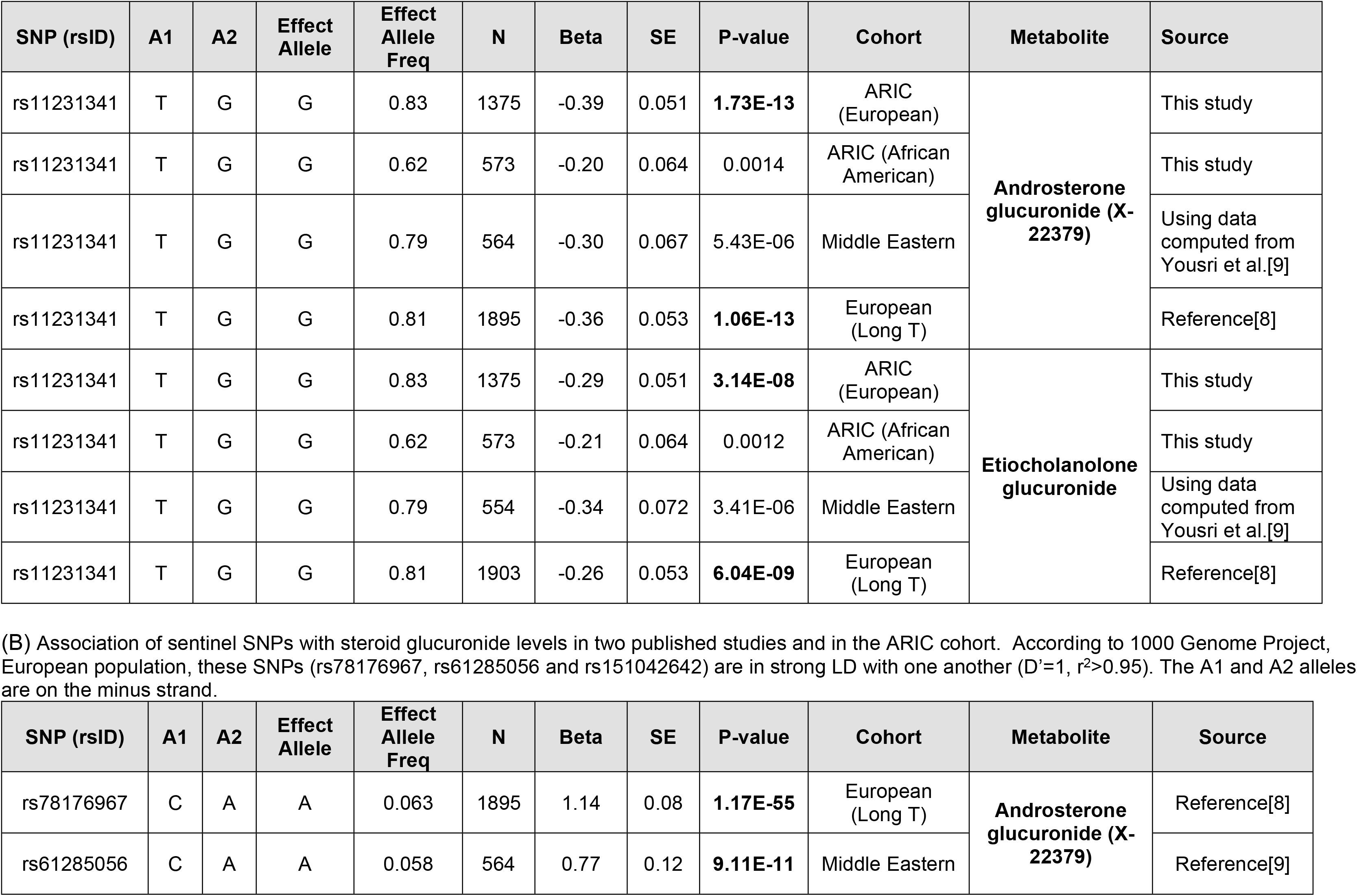
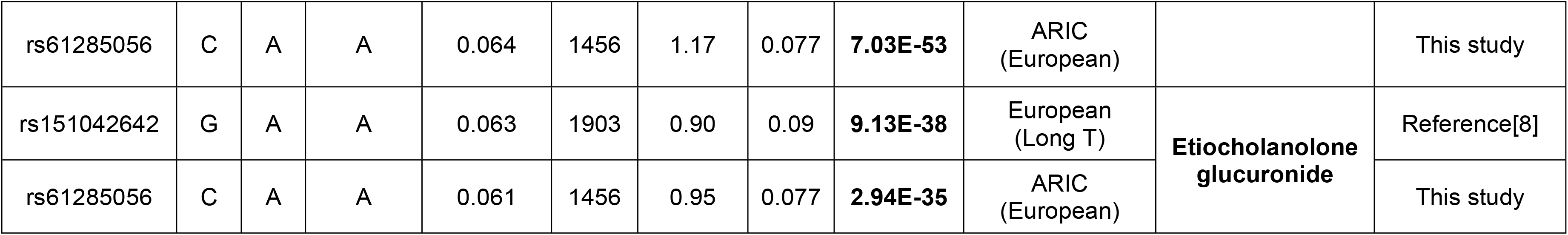
Association of (A) SLC22A24 p.Tyr501Ter (rs11231341, c.1503T>G) and (B) sentinel SNPs with two steroid conjugates in three cohorts. (A) Association of SLC22A24 p.Tyr501Ter (rs11231341, c.1503T>G) with steroid glucuronide levels. Major allele G (SLC22A24-501Ter) is associated with lower steroid glucuronide levels.

| SNP (rsID) | A1 | A2 | Effect Allele | Effect Allele Freq | N | Beta | SE | P-value | Cohort | Metabolite | Source |
| --- | --- | --- | --- | --- | --- | --- | --- | --- | --- | --- | --- |
| rs11231341 | T | G | G | 0.83 | 1375 | -0.39 | 0.051 | **1.73E-13** | ARIC (European) | **Androsterone glucuronide (X-22379)** | This study |
| rs11231341 | T | G | G | 0.62 | 573 | -0.20 | 0.064 | 0.0014 | ARIC (African American) |  | This study |
| rs11231341 | T | G | G | 0.79 | 564 | -0.30 | 0.067 | 5.43E-06 | Middle Eastern |  | Using data computed from Yousri et al.[9] |
| rs11231341 | T | G | G | 0.81 | 1895 | -0.36 | 0.053 | **1.06E-13** | European (Long T) |  | Reference[8] |
| rs11231341 | T | G | G | 0.83 | 1375 | -0.29 | 0.051 | **3.14E-08** | ARIC (European) | **Etiocholanolone glucuronide** | This study |
| rs11231341 | T | G | G | 0.62 | 573 | -0.21 | 0.064 | 0.0012 | ARIC (African American) |  | This study |
| rs11231341 | T | G | G | 0.79 | 554 | -0.34 | 0.072 | 3.41E-06 | Middle Eastern |  | Using data computed from Yousri et al.[9] |
| rs11231341 | T | G | G | 0.81 | 1903 | -0.26 | 0.053 | **6.04E-09** | European (Long T) |  | Reference[8] |

| SNP (rsID) | A1 | A2 | Effect Allele | Effect Allele Freq | N | Beta | SE | P-value | Cohort | Metabolite | Source |
| --- | --- | --- | --- | --- | --- | --- | --- | --- | --- | --- | --- |
| rs78176967 | C | A | A | 0.063 | 1895 | 1.14 | 0.08 | **1.17E-55** | European (Long T) | **Androsterone glucuronide (X-22379)** | Reference[8] |
| rs61285056 | C | A | A | 0.058 | 564 | 0.77 | 0.12 | **9.11E-11** | Middle Eastern |  | Reference[9] |
| rs61285056 | C | A | A | 0.064 | 1456 | 1.17 | 0.077 | **7.03E-53** | ARIC<br>(European) |  | This study |
| rs151042642 | G | A | A | 0.063 | 1903 | 0.90 | 0.09 | **9.13E-38** | European<br>(Long T) | **Etiocholanolone**<br>**glucuronide** | Reference[8] |
| rs61285056 | C | A | A | 0.061 | 1456 | 0.95 | 0.077 | **2.94E-35** | ARIC<br>(European) |  | This study |

### Phylogenetic analysis, comparative structure modeling, and metabolomic studies reveal SLC22A24 as an organic anion transporter

#### Phylogenetic analysis

First, we determined the relations between the human SLC22A24 and other human, mouse and rat SLC22 family members with known substrates (Fig 2A). In the phylogenetic analysis we also included genes within the locus between SLC22A8 and SLC22A9, known as the “unknown substrate transporters (USTs)”[31]. Notably, the number of genes annotated as USTs in mouse and rat is different from human. Mouse USTs include Slc22a26, a27, a28, a29, a30, a9/a19. Rat USTs include Slc22a9/a24, a9/a25, Ust4r, Ust5r. This analysis showed that human SLC22A24 lacks close human paralogs and rodent orthologs, unlike some well-characterized SLC22 family members such as SLC22A1 and SLC22A2 (Fig 2A).

**Fig 2.**
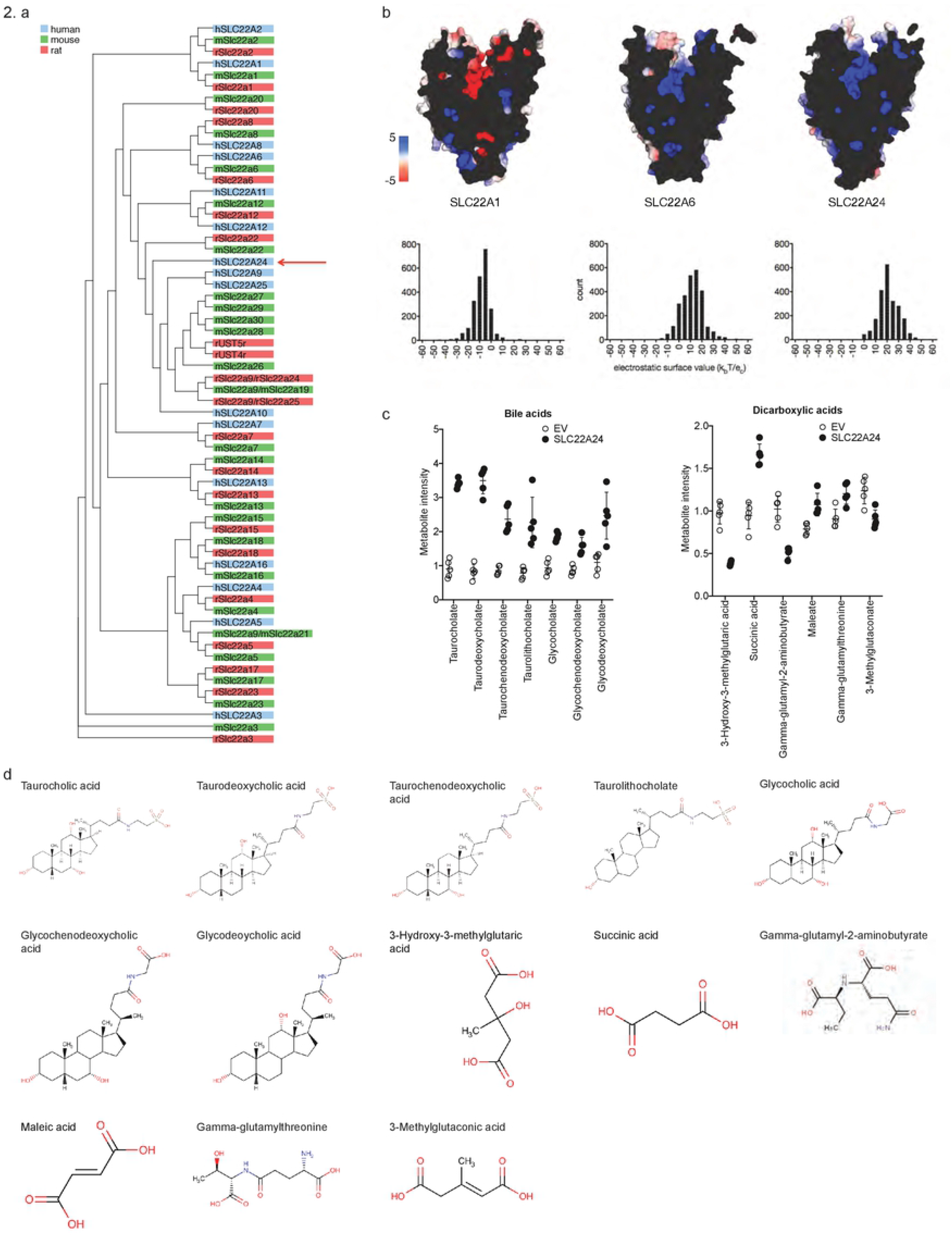
Phylogenetic tree analysis, comparative structure models and electrostatic potential analysis, and metabolomic study support the role of SLC22A24 as anion transporter. **A**) Sequence comparison of human, mouse, and rat protein sequences for all characterized SLC22 family members. Proteins are colored by organism. **B**) Top. Surface representations of the calculated electrostatic potential of the human SLC22A1, a characterized cationic transporter, the human SLC22A6, a characterized anionic transporter, and the human SLC22A24. The cross-section allows for visual inspection of the predicted binding pocket. Bottom. Calculated distribution of the electrostatic surface potential within 7Å of the center of the binding pocket for each of the transporters. **C)** Cellular metabolite levels are significantly different between HEK293 Flp-In cells transiently transfected with empty vector (EV) or *SLC22A24* for seven bile acids and six dicarboxylic acids among the top 50 metabolites. **D)** Chemical structures of the seven bile acids and six dicarboxylic acids.

#### Comparative structure modeling

Due to the lack of available crystal structures for most SLC transporters, we modeled human SLC22 members using the crystal structure of the human GLUT3 (SLC2A3) as a template. We hypothesized that despite the low sequence identity, the alignment would lead to an accurate assignment of residues along the helices, thus allowing an analysis of the electrostatic potential within the orthosteric site. As a control, we first modeled SLC22A1, a well-characterized organic cation transporter, and SLC22A6, an organic anion transporter (Fig 2B). A visual inspection and quantitative analysis of the electrostatic surface potential within the pocket was in agreement with the charges of known substrates. That is, the dominant charge of SLC22A1 was negative and SLC22A6 was positive, consistent with binding and translocation of cationic and anionic substrates, respectively. The extension of the same methodology to SLC22A24 showed the pocket to carry an overall positive charge, implying an affinity for anionic compounds.

#### Metabolomic study

Large-scale metabolomic studies have been previously used in our laboratory and others to identify transporter substrates in an unbiased manner[32–34]. The technological advances of state-of-the-art metabolomic methods have allowed for increased sensitivity and accuracy in the quantification of small molecules, elevating the number of detectable endogenous metabolites achieving a Tier1 standard for identification to over 700[35] (**S3 Table**). In this study, we transiently transfected HEK293 Flp-In cells with either empty vector (EV) or *SLC22A24*. *SLC22A1* was transiently transfected as positive control (**S3 Table**). Results in **S3 Table** show that many known metabolites of SLC22A1 were significantly different between SLC22A1 and EV transfected cells, for examples known substrates thiamine and acylcarnitines[32, 36]. A total of 50 metabolites reached the threshold of significance for difference in abundance between EV cells and SLC22A24 cells (Fig 2C-D). Among the top 10 metabolites, four were bile acids (taurocholic acid, taurodeoxycholic acid, glycocholic acid and taurochenodeoxycholic acid), which have steroid backbones, and two were dicarboxylic acids (3-hydroxy-3-methylglutarate and succinic acid) (Fig 2). Steroids and steroid conjugates were not above the threshold of detection in the cellular metabolomic analysis (**S3 Table**). Although most of the metabolites were present in greater abundance in the SLC22A24 expressing cells compared with EV cells, a few were present at lower abundance. Other anionic metabolites among the top 50 have monocarboxylic acid, phosphate, or sulfate moieties (**S3 Table**).

### A diverse set of *in vitro* transporter assays supports the role of SLC22A24 as organic anion transporter

We performed an uptake screen of 21 radiolabeled canonical substrates of various SLC22 family members, spanning cations, anions and zwitterions. The experiments confirmed that SLC22A24 is a transporter of select organic anions (**S4 Table**).

Cells transiently transfected with *SLC22A24* accumulated steroid anionic compounds, including [^3^H]-taurocholic acid (TA) (10-20 fold over vector control), [^3^H]-glycocholic acid (GA) (10-20 fold), [^3^H]-estrone sulfate (ES) (5-10 fold), [^3^H]-estradiol-17β-glucuronide (EG) (3-5 fold), and [^3^H]-androstanediol glucuronide (AG) (5-15 fold) (Fig 3A). The uptakes of [^3^H]-estrone sulfate, [^3^H]-estradiol-17β-D-glucuronide, [^3^H]-androstanediol glucuronide, [^3^H]-taurocholic acid and [^3^H]-glycocholic acid were time dependent, reaching plateaus at approximately 5 min. Kinetic parameters were evaluated at 1 min. The uptake kinetics of [^3^H]-estrone sulfate, [^3^H]-estradiol-17β-D-glucuronide, [^3^H]-taurocholic acid, and [^3^H]-glycocholic acid exhibited saturable characteristics at higher concentrations, with Km values (mean ± SD) of 8.6 ± 0.3 μM, 17.5 ± 0.2 μM, 10.5 ± 1.1 μM, and 33.4 ± 2.5 μM, respectively (Fig 3B, **S5 Table**). Similar values were observed for SLC22A8 (**S5 Table**). The kinetic parameters of androstanediol-3-glucuronide could not be determined accurately in SLC22A24 cells due to a solubility limit of 300 µM (**S2 Fig**). Using the plateau at 300 µM as an approximation, the Km of androstanediol glucuronide for SLC22A24 was estimated to be 741 ± 253 μM, which is higher than for SLC22A8.

**Fig 3.**
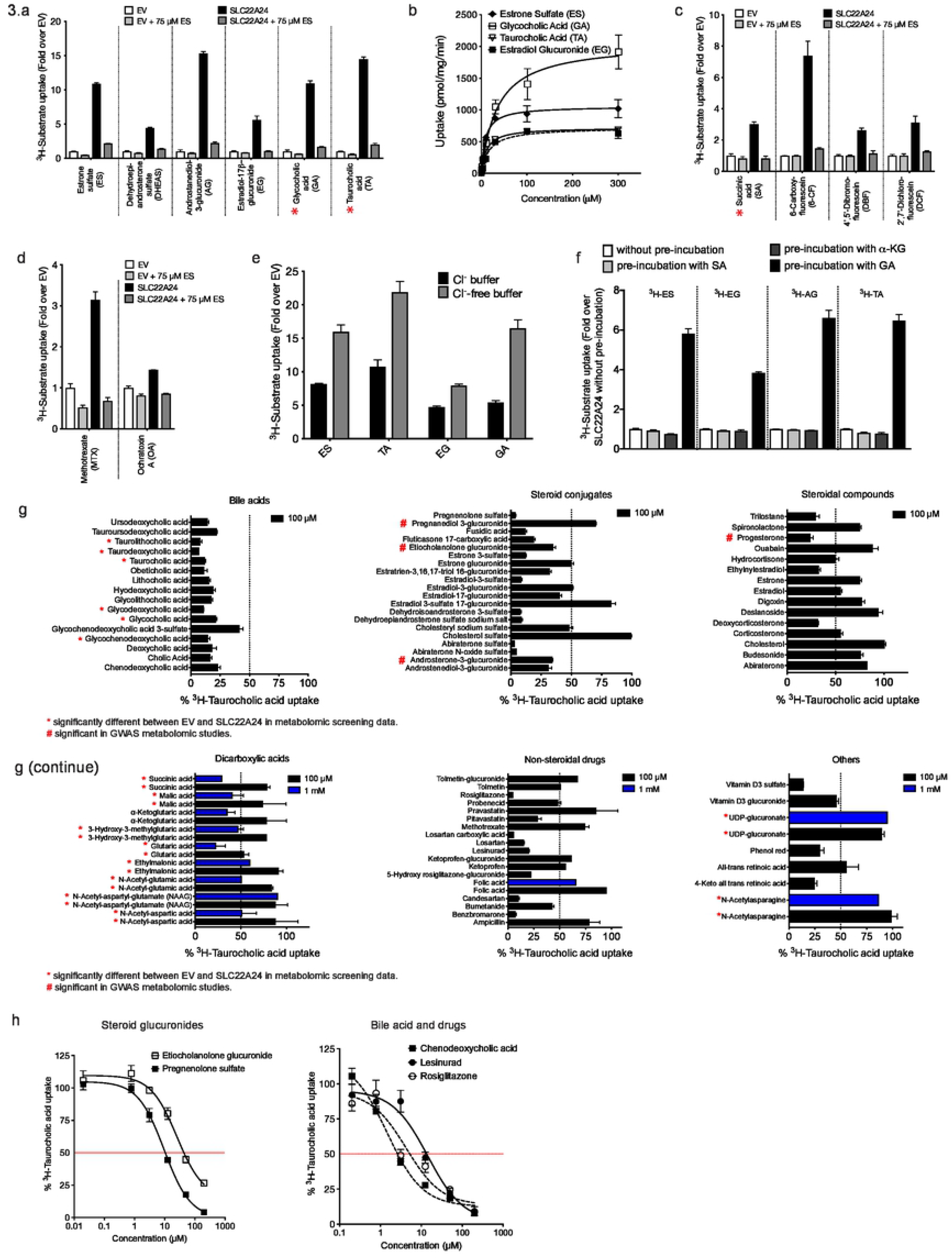
Uptake and inhibition studies of various anions in HEK293 Flp-In cells stably expressing SLC22A24. **A**) Uptake of steroid sulfate or glucuronide (ES, DHEAS, AG, EG) conjugates and bile acids (GA, TA). Uptake of these anions was significantly different between EV and *SLC22A24* transfected cells. Uptake was significantly reduced in the presence of excess unlabeled estrone sulfate. Figure shows a representative plot from one experiment (mean ± S.D. from three replicate wells). The experiments were repeated at least one time and showed similar results. **B**) Kinetics of uptake of four anions for SLC22A24. The uptake rate was evaluated at 2 minutes. Each point represents the uptake (mean ± S.D.) in the cells stably expressing SLC22A24, normalized to the empty vector control. The data were fit to a Michaelis−Menten equation. Figure shows a representative plot from one experiment. The mean and S.D. of the kinetic parameters from two experiments are shown in **S5 Table**. **C, D**) Uptake of anions with carboxylic acid and dicarboxylic acid groups. Uptake of these anions was significantly different between cells stably expressing SLC22A24 and the empty vector control cells. Uptake was reduced significantly in the presence of excess unlabeled estrone sulfate. Figure shows a representative plot from one experiment (mean ± S.D. from three replicate wells). The experiments were repeated at least one time and showed similar results. **E**) Uptake of four anions in HEK293 Flp-In cells stably expressing SLC22A24 with chloride-containing and chloride-free buffer. Each bar represents uptake (mean ± S.D. from three replicate wells) in the SLC22A24-transfected cells normalized to the empty vector control. The experiments were repeated at least one time and showed similar results. **F**) Trans-stimulation of SLC22A24–mediated estrone sulfate uptake. The uptake of estrone sulfate was trans-stimulated by separately preloading the cells with 2 mM of succinic acid, alpha-ketoglutrate acid, or glutaric acid for 2 h. After preloading, the uptake of the anions was measured after 10 min. Data are means ± S.D. The experiments were repeated one time, in triplicate, and show similar results. All experiments were normalized by setting the uptake of SLC22A24-expressing cells without preloading to 100%. **G**) Inhibition of [^3^H]-taurocholic acid uptake (with 1 µM unlabeled taurocholic acid) in cells stably expressing SLC22A24 by various bile acids, steroids, steroid conjugates, dicarboxylic acids, non-steroidal drugs, and other substances. All compounds were screened at 100 µM, and for some compounds, also at 1 mM. Uptake of [^3^H]-taurocholic acid was stopped after 10 min. Values represent the mean ± S.E.M. (from at least one experiment with three replicate wells each time). *Compounds that are statistically significant in the metabolomic analysis. ^#^Steroidal compounds that are statistically significant in the genome-wide association studies. **H**) Inhibition plots for SLC22A24-mediated taurocholic acid uptake in HEK293 Flp-In cells stably expressing SLC22A24. Cells were incubated with HBSS buffer containing taurocholic acid (1 µM) for 10 min with or without various concentrations of the compounds. Values are presented as mean ± S.D. of taurocholic acid uptake from three replicate wells determined in a single experiment. The experiments were repeated once and similar results were obtained. Representative curves of the SLC22A24-mediated taurocholic acid uptake inhibition by steroid conjugates (progesterone sulfate, etiocholanolone glucuronide), bile acid, and drugs (chenodeoxycholic acid, lesinurad, rosiglitazone). See Table 2 for the IC_50_ values.

Next, we characterized the uptake of monocarboxylic acid and dicarboxylic acid derivatives, which are known substrates of other anion transporters, including several prescription drugs. Among the compounds that were identified as substrates of SLC22A24, two were dicarboxylic acids (i.e. succinic acid and 6-carboxyfluorescein, 6-CF) and two were monocarboxylic acids (i.e. 4’,5’- dibromofluorescein, DBF, and 2’7’-dichlorofluorescein, DCF) (Fig 3C). However, other anion substrates of SLC22A6, SLC22A7 or SLC22A8, were determined not to be substrates of SLC22A24 including *para*-aminohippuric acid (PAH), uric acid, orotic acid, alpha-ketoglutaric acid and tetradecanedioic acid (**S4 Table**). The prescription drug, methotrexate, and the mycotoxin, ochratoxin A, showed significant uptake by SLC22A24 over EV cells (Fig 3D), whereas tenofovir and adefovir, which have phosphate moieties, were not substrates of the transporter (**S4 Table**). Further, none of the tested cationic and zwitterionic substrates of SLC22A1-SLC22A5 showed significant (>1.5 fold) accumulation in SLC22A24 expressing cells compared to control cells (**S4 Table**).

Collectively, the data indicate that SLC22A24 transports various organic anions, including steroid compounds with anionic moieties (sulfates, carboxylic acid, glucuronide) and non-steroid compounds with dicarboxylic acid moieties.

### SLC22A24 transport is chloride sensitive and transtimulated by glutaric acid

We examined the underlying transport mechanism of SLC22A24 in order to determine whether it relies on facilitative diffusion or on secondary active transport of ions. Uptake of [^3^H]-estrone sulfate was not affected by pH between 5.5 to 8.5 and was not dependent on the presence of Na^+^ ions (**S3 Fig**). However, uptake of estrone sulfate, as well as other steroid conjugates, was significantly affected by the presence of Cl^−^ ions. Cl^−^-free buffer nearly doubled the uptake of all tested compounds (Fig 3E). In addition, we tested trans-stimulation of SLC22A24-mediated uptake by preloading the cells with succinic acid, alpha-ketoglutaric acid, and glutaric acid, which are known to trans-stimulate other organic anion transporters[37–39]. Among the three, only glutaric acid significantly increased the uptake of [^3^H]-estrone sulfate, [^3^H]-estradiol glucuronide, [^3^H]-androstenediol glucuronide and [^3^H]-taurocholic acid (Fig 3F).

### SLC22A24 is inhibited by anionic compounds from different chemical classes

The goal of the inhibition studies was to confirm that solutes identified in the metabolomic studies were ligands of SLC22A24 and to identify additional ligands of the transporter. A second goal was to determine whether known inhibitors of other organic anion transporters in the SLC22 family were inhibitors of SLC22A24. A total of 85 compounds were selected for inhibition studies of SLC22A24-mediated [^3^H]-taurocholic acid uptake (**S6 Table**, Fig 3G). These included

- 4 steroid metabolites identified in GWAS at p<1×10^−6^
- 20 known inhibitors or substrates of other organic anion transporters
- 17 steroids or bile acids which are used clinically
- 8 dicarboxylic acid metabolites identified in our metabolomic study
- 20 steroid conjugates

At 100 µM, 43 of the 86 test-compounds inhibited the SLC22A24-mediated [^3^H]-taurocholic acid uptake by >50%. All 16 bile acids, 5 of the 15 unconjugated steroids, 15 of the 20 steroid conjugates, and 12 of the 22 other anionic non-steroidal chemical compounds inhibited taurocholic acid uptake by >50% (Fig 3G, **S6 Table**). Many sulfate conjugates including sulfate conjugates of steroids inhibited taurocholic acid uptake. Some non-steroidal compounds known to inhibit organic anion transporters also inhibited SLC22A24-mediated taurocholic acid uptake potently (e.g. losartan, rosiglitazone, lesinurad, benzbromarone) or partially (e.g. probenecid, bumetanide, ketoprofen). Three steroid metabolites identified in the GWAS, progesterone, androsterone glucuronide, and etiocholanolone glucuronide inhibited taurocholic acid uptake >50%. Consistent with these results, SNPs in SLC22A24 were associated with progesterone, androsterone glucuronide and etiocholanolone glucuronide at genome-wide significance levels (p<5×10^−8^), whereas associations were weaker for pregnanediol-3-glucuronoide (p=2×10^−7^), which was a weaker inhibitor of SLC22A24-mediated taurocholic acid uptake (Fig 3G)[8]. At a high concentration (1 mM), five of the nine dicarboxylic acids inhibited taurocholic acid uptake by approximately 50% (Fig 3G). However, UDP-glucuronate and N-acetylasparagine did not inhibit taurocholic acid uptake even at 1 mM.

Ten compounds were selected for further IC_50_ studies (Table 2). We specifically focused on the steroids and steroid conjugates reported in GWAS and the steroid conjugates with the strongest inhibition in the initial screen. Two bile acids, chenodeoxycholic acid and ursodeoxycholic acid, were selected because they are used clinically to treat gallstone and related cholestatic liver disease [40, 41]. Two non-steroidal drugs, rosiglitazone and lesinurad, were selected because of their strong inhibition in the initial screen. Pregnenolone sulfate, chenodeoxycholic acid, and rosiglitazone exhibited potent IC_50_ values (IC_50_ < 5 µM). Progesterone, ursodeoxycholic acid, and lesinurad were also inhibitors with IC_50_ < 20 µM. Finally, the two steroid glucuronides, etiocholanolone glucuronide and androsternediol-3-glucuronide, achieved IC_50_ values of 29±5 μM and 21±11 μM, respectively (Table 2, Fig 3H).

**Table 2.**
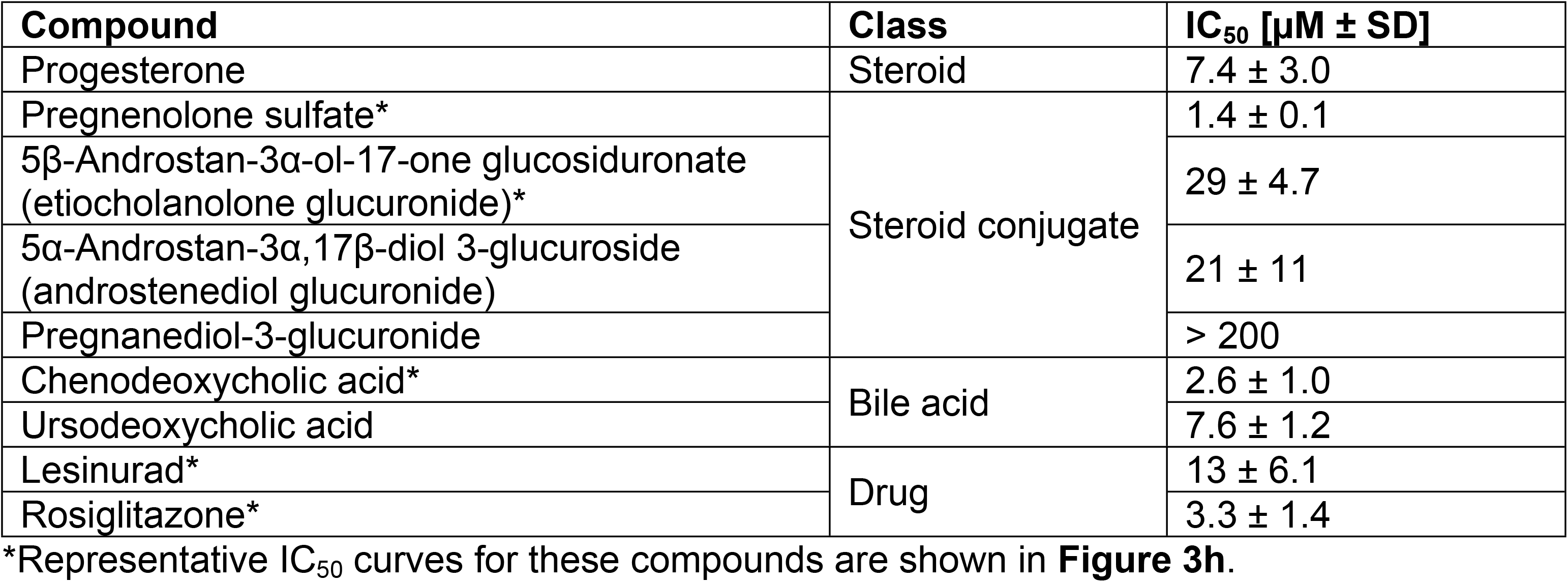
Inhibition potencies of steroid, steroid conjugates, bile acids and drugs for SLC22A24-mediated taurocholic acid uptake. The experiments were repeated once to obtain IC_50_ mean ± SD (the inhibition concentration at 50% taurocholic acid uptake).

| Compound | Class | IC<sub>50</sub> [μM ± SD] |
| --- | --- | --- |
| Progesterone | Steroid | 7.4 ± 3.0 |
| Pregnenolone sulfate* | Steroid conjugate | 1.4 ± 0.1 |
| 5β-Androstan-3α-ol-17-one glucosiduronate (etiocholanolone glucuronide)* |  | 29 ± 4.7 |
| 5α-Androstan-3α,17β-diol 3-glucuroside (androstenediol glucuronide) |  | 21 ± 11 |
| Pregnanediol-3-glucuronide |  | > 200 |
| Chenodeoxycholic acid* | Bile acid | 2.6 ± 1.0 |
| Ursodeoxycholic acid |  | 7.6 ± 1.2 |
| Lesinurad* | Drug | 13 ± 6.1 |
| Rosiglitazone* |  | 3.3 ± 1.4 |

\*Representative IC<sub>50</sub> curves for these compounds are shown in **Figure 3h**.

### Species differences in the substrate selectivity of SLC22A24 orthologs exist

Data from Ensembl[42, 43] revealed that SLC22A24 1:1 orthologs were present in some, but not all mammals and were absent in non-mammalian vertebrates (e.g., fish, reptiles and birds). Fig 4A shows the phylogenetic analysis of ten SLC22A24 1:1 orthologs in mammals along with other relevant organic anion transporters in the SLC22 family (see also **S4 Fig**). Included in the figure are SLC22A24 orthologs from apes (orangutan, chimpanzee, gorilla and gibbon), lemur (mouse lemur and coquerels sifaka), rabbit, and horse. We compared the function of the human SLC22A24 with ortholog representatives from the different branches of the phylogenic tree (Fig 4B), that of the horse, mouse lemur, and rabbit. We also included the rat Slc22a9/Slc22a24, previously known as Slc22a19 and Slc22a9, but most recently assigned the gene name *Slc22a24* [44]. All five species (human, horse, mouse lemur, rabbit, and rat) showed significant uptake of estrone sulfate (>2.5 fold) compared to empty vector transfected cells (Fig 4B). There were substrate specificity differences between the human SLC22A24 and the orthologs of the other species investigated. Notably:

- Horse SLC22A24 significantly transports taurocholic acid (∼2 fold), estrone sulfate (∼3 fold), and estradiol glucuronide (∼2 fold), but not androstenediol glucuronide (∼1.2 fold).
- Mouse lemur SLC22A24 transports estrone sulfate (∼17 fold), androstenediol glucuronide (∼2 fold) and estradiol glucuronide (∼11 fold) but not taurocholic acid.
- Rabbit SLC22A24 significantly transports taurocholic acid (∼11 fold), estrone sulfate (∼5 fold), androstenediol glucuronide (∼4 fold), and estradiol glucuronide (∼2 fold).
- Rat Slc22a9/Slc22a24, which has the lowest sequence similarity to human protein out of the three species above, does not significantly transport taurocholic acid or androstenediol glucuronide.

**Fig 4.**
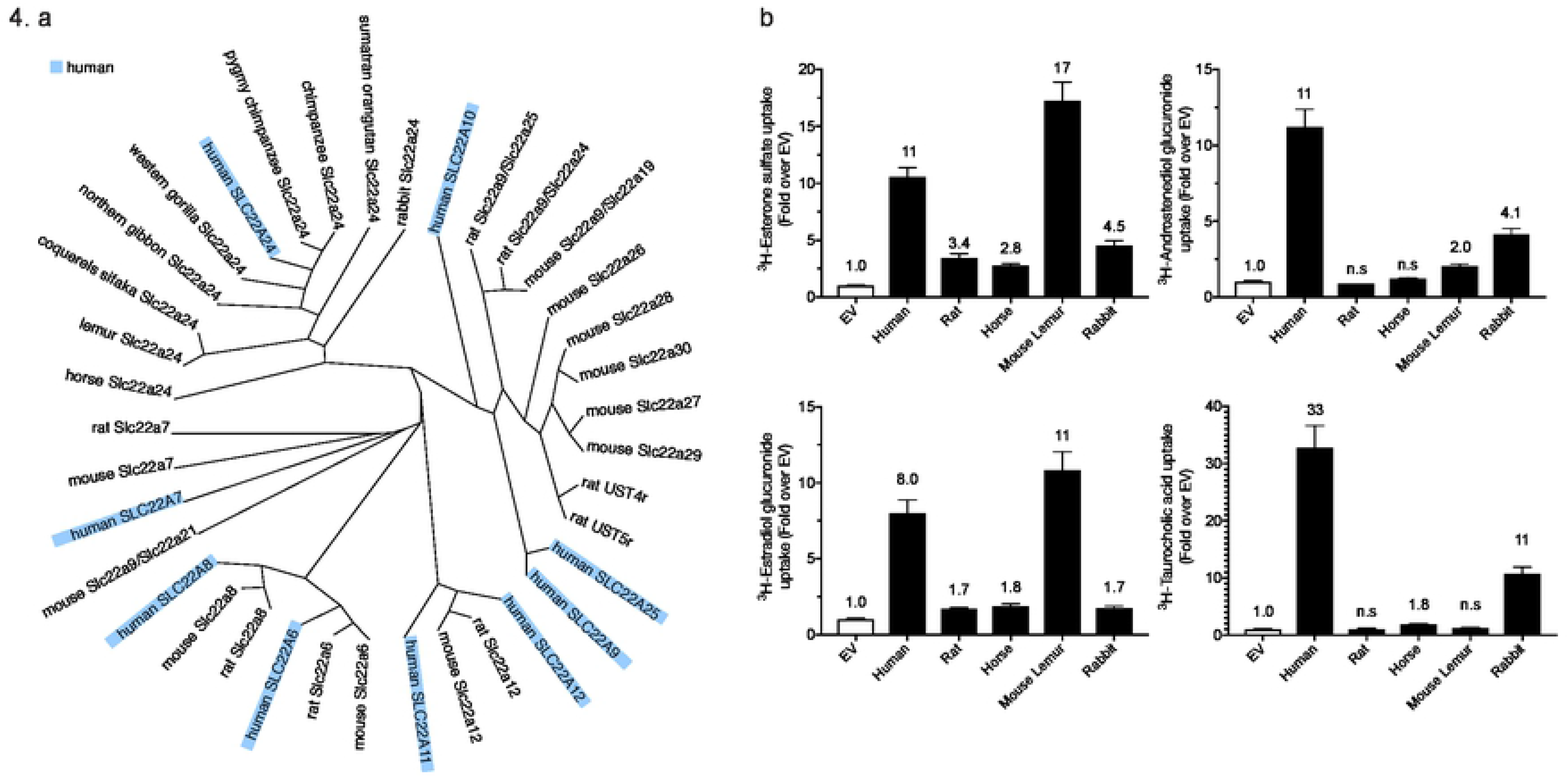
Phylogenetic tree analysis and uptake of anions by species orthologs of SLC22A24. **A**) Sequence comparison of characterized human anion transporters and select mammalian orthologs. **B**) Fold uptake of four substrates by species orthologs of SLC22A24 and rat Slc22a9/Slc22a24 over empty vector control. Uptake of esterone sulfate, androstenediol glucuronide, estradiol glucuronide, and taurocholic acid was performed in HEK293 Flp-In cells transiently expressing *SLC22A24* from human, mouse lemur, rabbit and horse or *Slc22a9/a24* from rat. Uptakes were performed for 10 minutes. The number above the bar is fold uptake over EV cells from one experiment (mean ± S.D., from four replicate wells). The experiments were repeated one time and similar results were obtained.

We also determined the plasma membrane expression of each of the above orthologs in HEK293 cells using a GFP-tag to ensure the proper localization of the expressed proteins (**S5 Fig**). All five orthologs were expressed on the plasma membrane, but the expression levels were variable. More pronounced plasma membrane expression was observed for the rabbit and human orthologs. The horse, rat, and the mouse lemur orthologs, were also present on the plasma membrane, but had a more diffuse intracellular expression (**S5 Fig**).

### SLC22A24 isoforms exhibit no significant differences in their substrate selectivity, while SLC22A24 p.Tyr501Ter variant exhibits no protein expression or transport function

According to UniProt, Ensembl, and NCBI RefSeq, the SLC22A24 transcript is identified in three major isoforms. Specifically, ENST00000612278.4 (552 amino acids, NM_001136506, A0A087WWM3), identified in this paper as the reference SLC22A24, ENST00000417740.5 (551 amino acids, no known NCBI reference, C9JC66), identified as SLC22A24-551, and ENST00000326192.5 (322 amino acids, NM_173586, Q8N4F4), identified as SLC22A24-322. The short SLC22A24-322 isoform was omitted from further analysis due to its lack of transport activity, expression from GFP fusion construct, and absence of key structural motifs (see **S6 Fig**). Transcripts of the two isoforms (SLC22A24 and SLC22A24-551) were detected using PCR and the sequences were confirmed. The SLC22A24-551 isoform exists due to the presence of two potential AG acceptor sites at the splice junction between exons 9 and 10, (indicated as position “1” or “2” on **S7A Fig)**, thus resulting in two different splice variants. The splice variants differ in two amino acids in position 533 and 534 (**S7B Fig**). Functional characterization of SLC22A24 and SLC22A24-551 indicated no significant differences in their uptake function. In particular, a similar fold-uptake of the two isoforms over control was observed for all representative steroid glucuronides, steroid sulfates, bile acids, methotrexate and succinic acid (Fig 5A).

**Fig 5.**
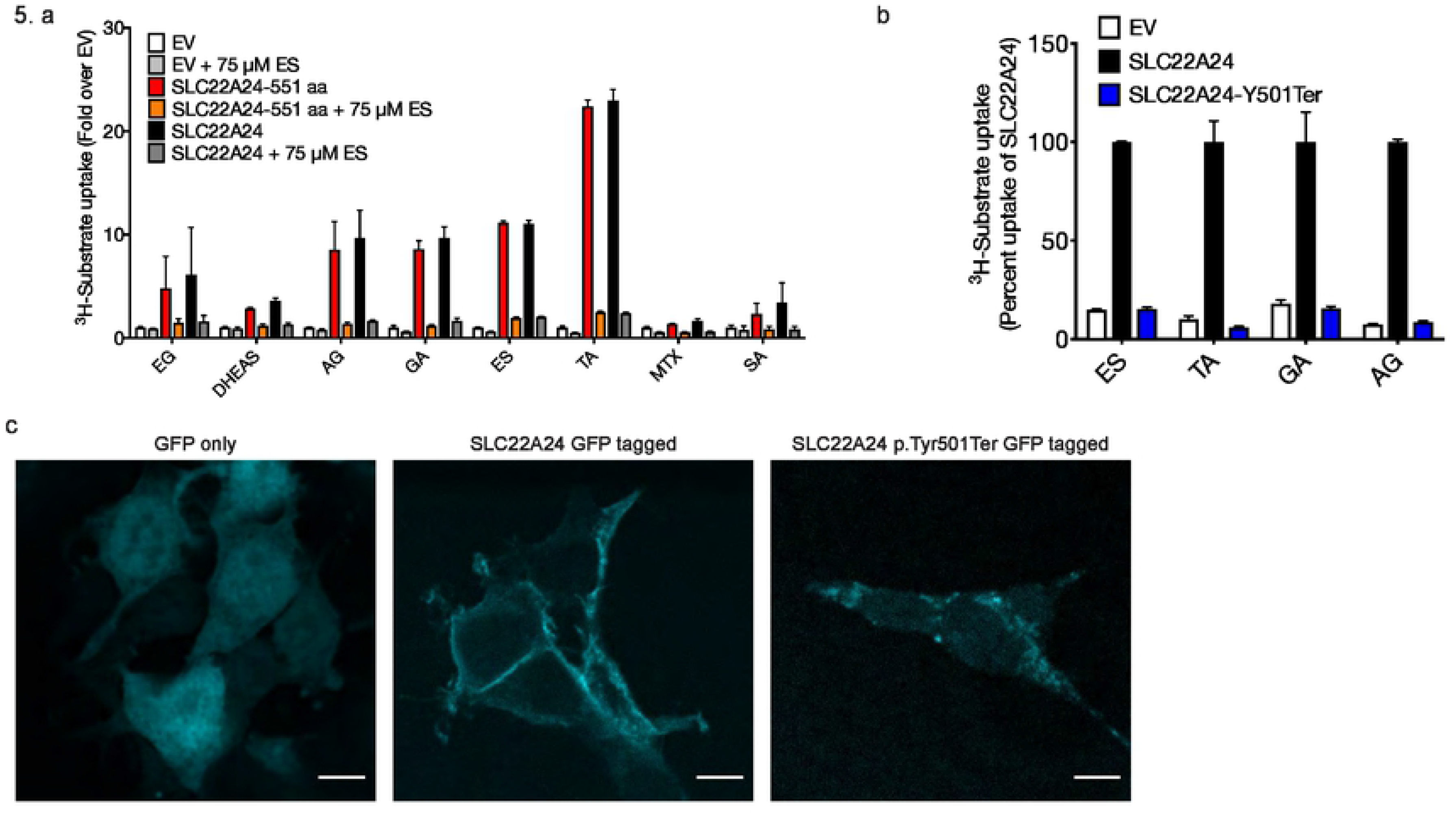
Uptake of various anions by two SLC22A24 isoforms and the effect of a common nonsense variant, p.Tyr501Ter on uptake and plasma membrane localization. **A**) Fold uptake over empty vector for different substrates. HEK293 Flp-In cells were transiently transfected with different constructs containing *SLC22A24* or *SLC22A24-551*. The experiments were repeated at least one time. Data represents means ± S.D. of two independent experiments. **B**) Transport activity for different substrates was compared in HEK293 Flp-In cells transiently transfected with *SLC22A24* or *SLC22A24* p.Tyr501Ter. The uptake of SLC22A24-expressing cells was set to 100%. Data represents means ± S.D. from three wells. The experiments were repeated one time and similar results were obtained. **C**) Membrane localization of SLC22A24 and SLC22A24-Y501Ter. HEK293 cells were transiently transfected with GFP fusion constructs and visualized by confocal microscopy.

Next, we characterized the nonsense variant in SLC22A24, which is associated at genome-wide significance levels with androsterone glucuronide and etiocholanolone glucuronide (Fig 1). A survey of the 1000 Genomes Project Phase 3, revealed that the G-allele of rs11231341, coding for p.Tyr501Ter, is the major allele, ranging from 51% in Mende in Sierra Leone to 92% in Peruvian in Lima, Peru [45]. In addition, the G-allele is the major allele (80%) in individuals from the Greater Middle East (GME) (The GME Variome Project, http://igm.ucsd.edu/gme/). Cells transiently transfected with SLC22A24 p.Tyr501Ter exhibited no transport function (Fig 5B). Consistent with the uptake studies, GFP-tagged SLC22A24 localized to the membrane, while GFP-tagged SLC22A24 p.Tyr501Ter did not (Fig 5C). Further, plasma membrane protein expression was evaluated using western blots for HEK293 Flp-In cells stably expressing SLC22A24 or transiently transfected with either empty vector only (EV, pCMV6-Entry), SLC22A24, or SLC22A24 p.Tyr501Ter (**S8A Fig**). All samples expressed the positive control, Na^+^/K^+^-ATPase. SLC22A24 was expressed in both the stable cell sample as well as the transiently transfected cells. SLC22A24 p.Tyr501Ter was not detected on the plasma membrane of the transiently transfected cells, despite having detectable mRNA levels (**S8B Fig**).

### Transcriptomic and proteomic studies reveal low and variable expression levels of multiple isoforms of SLC22A24

#### Transcriptomic studies

PCR identified the transcripts to be expressed at variable levels in the 14 tissues tested (pooled from various samples, see **S7 Table**), in addition to kidney and liver (Fig 6A). SLC22A24 and SLC22A24-551 were found at moderate levels in nine tissues including total brain, liver, skeletal muscle, kidney, pancreas, colon, spleen, testis, and thymus (Fig 6A). Interestingly, the Kidney Interactive Transcriptomics database, which is a repository for single cell RNAseq data from human kidney, shows SLC22A24 to be expressed specifically in the proximal tubule segment 3 (S3), and not in other regions of the human kidney (http://humphreyslab.com/SingleCell/search.php, Fig 6B)[46], providing support for low overall transcriptomic expression in the kidney.

**Fig 6.**
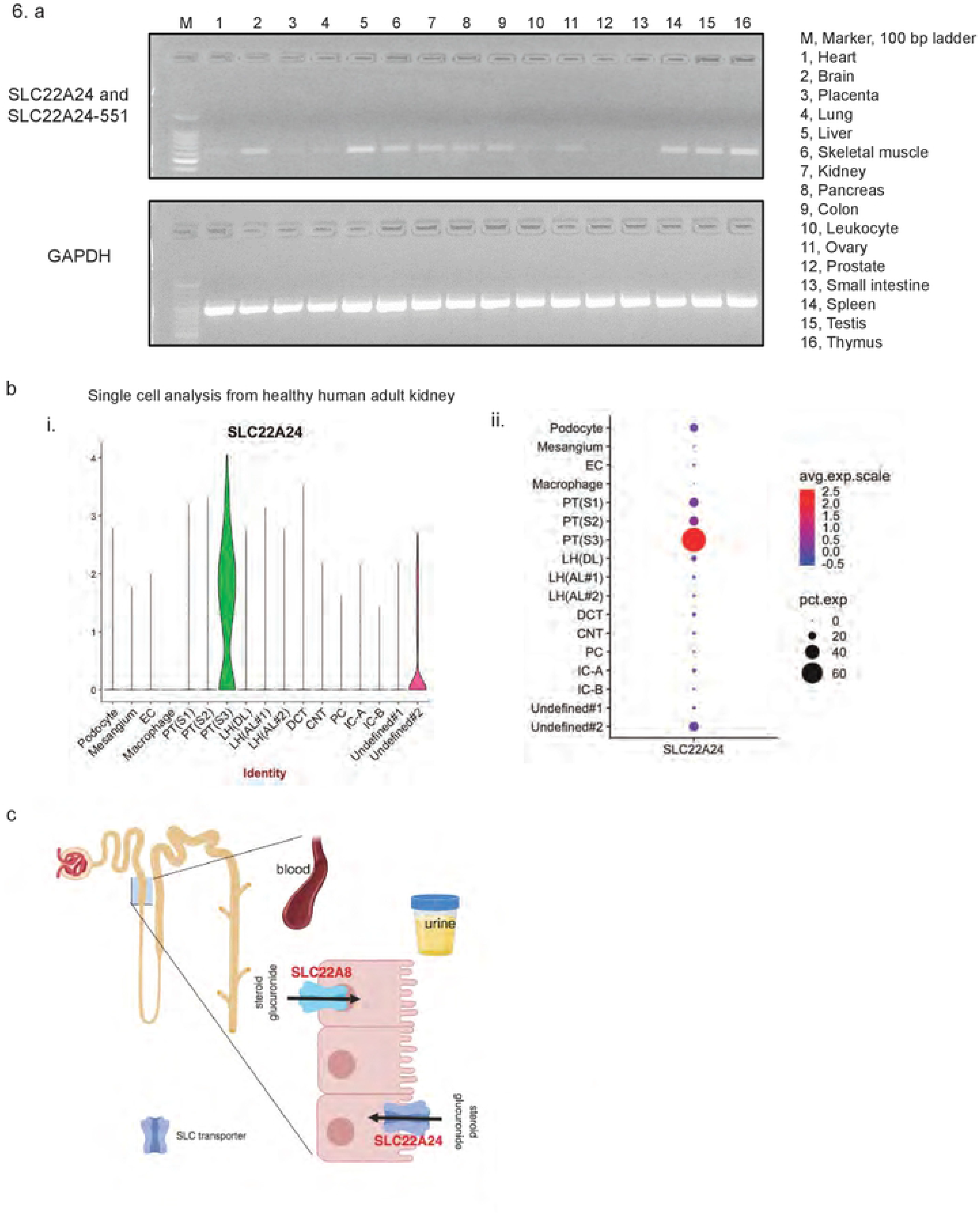
Expression of *SLC22A24* mRNA transcripts in human tissues and cells, and a proposed model for its functional role in the disposition of steroid conjugates in the kidney. **A**) Expression of *SLC22A24* mRNA in various pooled samples was detected by RT-PCR. Forty and 30 cycles were used for amplifying *SLC22A24* and *GAPDH* mRNA, respectively. **B**) *SLC22A24* expression from human kidney single cell datasets (Healthy Human Adult Kidney (Complete: 4,524 nuclei)) were obtained from Kidney Interactive Transcriptomics (http://humphreyslab.com/SingleCell/). Two different plots (i and ii) show the regions of the kidney and the abundance of *SLC22A24* mRNA transcripts. EC, endothelial cell; PT, proximal tubule; LH(AL), loop of Henle ascending loop; LH(DL), loop of Henle descending loop; DCT, distal convoluted tubule; CNT, connecting tubule; PC, collecting duct-principal cell; IC-A, intercalated cell type A; IC-B, intercalated cell type B. Overall, the plots indicate that *SLC22A24* is selectively expressed in the proximal tubule segment 3 (PT(S3)). **C**) Proposed mechanism of steroid glucuronide handling in the kidney by SLC22 transporters. The scheme illustrates that SLC22A24 is on the apical membrane of the proximal tubule and functions in the reabsorption of steroid glucuronides. Conversely, SLC22A8, localized to the basolateral membrane, functions in the secretion of steroid glucuronides[119]. This scheme was created with BioRender (https://biorender.com/).

#### Proteomic studies

Using LC-MS/MS Selective Reaction Monitoring (SRM) approach, we evaluated protein levels of the two SLC22A24 isoforms in 20 renal cortical samples from self-reported African Americans (age [mean ± SD]: 13 ± 8 years; 50% males). SRM is a sensitive mass spectrometry technique that is capable of accurate protein detection and quantification[47]. Three peptides were designed to detect and quantify the protein abundance (**S9 Fig**). Peptide 1 detected both SLC22A24 and SLC22A24-551, while peptides 2 and 3 were specific to SLC22A24 and SLC22A24-551, respectively. Surprisingly, isoforms SLC22A24 and SLC22A24-551 exhibited a mutually exclusive expression pattern (**S9 Table**), that is, each individual sample expressed only one of the two isoforms. Genotyping 20 renal cortical samples from African Americans, we identified the c.1503-G allele to have an allele frequency of 67.5% (Hardy-Weinberg p-value = 0.4), consistent with known allele frequencies in publicly available sources. Ten c.1503-GG, seven c.1503-GT, and three c.1503-TT were used for the SRM analysis (*SLC22A24* c.1503-GG codes for SLC22A24 p.Tyr501Ter homozygotes). In line with the stop codon at amino acid position 501, the SRM analysis showed that the renal cortical tissues with the genotype c.1503-GG have no detectable protein abundance using peptides 2 and 3 (**S9 Table**).

## Discussion

Steroids including sex steroids and glucocorticosteroids have pleiotropic actions in reproductive and other tissues in the body and impact numerous biological processes[1, 48, 49]. In particular, they influence body composition, glucose and lipid metabolism, cardiovascular and immunological functions, and reproductive fitness[49]. Accordingly, there is a vast literature focused on the study of the action, regulation, metabolism and disposition of steroids[1]. GWAS have revealed a number of genes that are involved in steroid metabolism and hormone binding as the major determinants of interindividual variation in the levels of many steroids and steroid conjugates [2–7, 50]. Intrigued by the association of a novel locus and an orphan transporter, SLC22A24, with metabolite levels in metabolomic GWAS, we performed experimental studies along with computational analysis to functionally characterize the transporter and further understand its physiological role. Our studies led to four major findings. First, SLC22A24 is an organic anion transporter, consistent with multiple GWAS focused on metabolites, and validated through comparative structure models, cell-based metabolomic screens, and functional studies. Second, SLC22A24 is conserved among higher order primates, but has diverged within lower level primates and other mammals. Third, two functional isoforms of SLC22A24 are detectable in the human kidney, and the transporter has a highly specific expression pattern in the nephron, localized to segment 3 of the proximal tubule. Fourth, the transporter has a common nonsense variant, p.Tyr501Ter (rs11231341), which is found at high allele frequencies in human populations, and is associated with low steroid and steroid conjugate levels (p<5×10^−8^). Below we discuss each of the major findings in detail.

The first major finding of the study is that SLC22A24 is an organic anion transporter (Fig 2 and Fig 3). The SLC22 family consists of organic ion transport proteins that are selective for molecules based on their charge. In general, family members have been grouped into organic anion, organic cation, and zwitterion transporters. Our first goal in discovering the functional role of SLC22A24 was to determine the charge specificity of the transporter. Based on results the GWAS (Fig 1) and comparative structure modeling (Fig 2), we hypothesized and validated in our *in vitro* and cellular metabolomic studies that SLC22A24 preferentially transports organic anions. However, it does not transport a number of organic anions transported by other organic anion transporters in the SLC22 family (SLC22A6, SLC22A7 and SLC22A8). For example, we did not detect transport of uric acid, tenofovir, cGMP, or alpha-ketoglutaric acid, which are canonical substrates of SLC22 organic anion transporters (**S5 Table**)[20, 51]. Further, our results showed that SLC22A24 is Cl^−^-dependent (Fig 3E). A similar chloride dependency has been observed for URAT1 (SLC22A12), a major uric acid reabsorptive transporter in the kidney[52], and OAT4 (SLC22A11), which uses the downhill flux of chloride to move its organic anion substrates across the apical membrane of the renal proximal tubule[37, 53]. Similar to OAT1 (SLC22A6), OAT3 (SLC22A8) and OAT4 (SLC22A11), SLC22A24 was also trans-stimulated by glutaric acid [37, 38, 54, 55], but not significantly by succinic acid and alpha-ketoglutaric acid (Fig 3F). In contrast, OAT1 and OAT3 are trans-stimulated by alpha-ketoglutaric acid as well as glutarate, consistent with a broader selectivity of these two transporters in comparison to SLC22A24.

Our second finding is that many species, including lower level primates and rodents, do not have orthologs of SLC22A24. Phylogenetic analysis reveals the presence of SLC22A24 orthologs in apes, lemur, and certain other mammalian species, but none in rodents. Human *SLC22A24*, located on chromosome 11q12, falls within a cluster of other SLC22 family genes called the Unknown Substrate Transporters (USTs), consisting of *SLC22A9*, *SLC22A10*, and *SLC22A25*. Sequence comparison indicates that human SLC22A24 is distant from those in mice and rats (mouse USTs include Slc22a26, a27, a28, a29, a30, a9/a19; rat USTs include Slc22a9/a24, a9/a25, Ust4r, Ust5r) (Fig 4A, **S4 Fig**). Similarly, the Ensembl database (version 95) identifies only 11 species as having direct orthologs of the human SLC22A24 (see ref. [43]). Previous studies have characterized the function of human liver specific *SLC22A9* (OAT7)[25], several rodent USTs, including the rat kidney Slc22a9/a24 (Oat5)[56], rat kidney Slc22a9/a25 (Ustr1/Oat8)[57, 58], mouse kidney Slc22a9/a19 (Oat5)[59–61], and mouse liver and kidney Slc22a30 (Oat9)[62]. All of the above transporters transport steroid sulfate conjugates and some other anions, such as ochratoxin A, but do not transport and are not inhibited by steroid glucuronide conjugates, which are substrates and inhibitors of the human SLC22A24[56, 58]. Consistent with these results, steroid glucuronides are not detectable in rat urine but are detectable in human urine[63]. Further, glucuronidated steroids are not detected in serum samples from rodents[64].

These studies suggest that in comparison to humans, rodents may have distinct conjugation pathways for steroids, e.g., sulfation, and rely on other transporters, perhaps SLC members outside of the SLC22 family, to transport steroids and steroid conjugates. Species differences in conjugation pathways and enzymes have been well described[65, 66]. For example, beta-estradiol is glucuronidated by different uridine 5’-diphosphate-glucuronosyltransferase (UGT) enzymes in rats and humans. These enzymes differ in their expression pattern and levels among various species[65]. Thus, it is not surprising that there are significant species differences in the expression pattern and levels of the proteins involved in the transport of steroids and their conjugates.

Further, even species that possess direct orthologs of SLC22A24 appear to have unique differences in their specificity for steroid glucuronides and bile acids. For example, the lemur ortholog of SLC22A24 transports estrone sulfate and various steroid glucuronides, but not taurocholic acid, which the human SLC22A24 transports, in addition to the steroid conjugates (Fig 4B). Although there are documented species differences in levels of bile salts and cholesterol[67, 68], it is difficult to speculate on the specific reasons that may be responsible for these differences. Future studies are needed to understand species differences in the pathways involved in steroid synthesis, metabolism, and disposition.

Our third finding is that the functional isoforms of the human SLC22A24, are expressed at lower levels in kidney and liver tissues compared to other known kidney-specific anion transporters from the SLC22 family, e.g., SLC22A6, SLC22A8, SLC22A11, and SLC22A12, as noted from the RNA-seq data generated by the Human Protein Atlas (HPA)[69] and the Genotype-Tissue Expression (GTEx) consortium (see **S10 Table**). Though expressed at lower levels, the primary finding that genetic variants in SLC22A24 associate with plasma levels of several metabolites (**S1 Table**), suggests that the transporter plays a critical and specific role in determining systemic levels of metabolites. The narrower substrate selectivity of SLC22A24 as compare to other SLC22 organic anion transporters, SLC22A6 and SLC22A8, which are broadly selective, further supports a specific role for the transporter in the kidney. The search results from the Kidney Interactive Transcriptomics database (http://humphreyslab.com/SingleCell/search.php), which shows SLC22A24 to be expressed specifically in S3 and not in the other regions of the human kidney suggest that SLC22A24 plays a specific role in the kidney in comparison with other renal organic anion transporters in the SLC22 family. The selective localization of SLC22A24 in the human kidney is consistent with the renal localization of rodent USTs, which are expressed in particular sub-regions of the kidney [56, 58, 60]. For example, rat Slc22a9 is expressed on the apical membrane of the collecting duct [58], and mouse Slc22a9/a19 is expressed on the apical membrane of the proximal tubules of the outer medulla [56, 60]. Collectively, the data suggest a specific role for SLC22A24 in the kidney, reminiscent of USTs in rodents, which do not express SLC22A24.

Our fourth finding is that the transporter has a common nonsense variant, p.Tyr501Ter (rs11231341), which is found at high allele frequencies in human populations, and is a major determinant of inter-individual variation in plasma levels of certain steroid and steroid conjugates (p<5×10^−8^). Premature stop codons sometimes trigger nonsense-mediated decay (NMD), which in this case might cause a rapid degradation of the SLC22A24 transcripts and result in the observed lack of detectable protein signal in the kidney samples from individuals who are homozygous for p.Tyr501Ter (**S8 Table**). The overall low mRNA transcript levels of SLC22A24 in publicly available databases (which either average expression levels from multiple samples or report results of pooled samples) may be due to the high allele frequency of the rs11231341 encoding the p.Tyr501Ter. Surprisingly, the premature stop codon has not been found in primates[70] (61 chimpanzee, 10 bonobo, 43 gorilla, 9 orangutan). However, according to the three ancient human genomes, the Altai Neandertal, Denisova, Ust-Ishim and Vindija, the ancient humans were either p.Tyr501Ter homozygous or heterozygous (available from the ancient genome browser, https://bioinf.eva.mpg.de/jbrowse/)[71, 72]. These data suggest that the mutation was introduced in the ancient human period, the reason for which remains an open evolutionary question.

Examination of GWAS and publicly available databases (GWAS Catalog, NCBI Phenotype-Genotype Integrator) revealed strong associations between SNPs in SLC22A24 and plasma levels of various metabolites, particularly those of steroid glucuronide conjugates (**S1 Table**), as well as steroid sulfate conjugates (**S1 Table**[25]). Although *in vitro* studies showed that SLC22A24 robustly transports bile acids, scanning across published GWAS did not reveal strong associations with bile acids (**S1 Table**). The direction of the association with the reduced function allele provides a viable hypothesis for the physiologic role of the transporter as a reabsorptive transporter in the kidney. SLC22A24 p.Tyr501Ter (rs11231341), which has no transport activity and no protein expression on the plasma membrane (Fig 5), is associated with lower plasma levels of steroid glucuronides in three independent cohorts and in multiple ethnic groups (Table 1). Thus, it is reasonable to speculate that individuals harboring p.Tyr501Ter would have reduced plasma steroid glucuronide levels due to their reduced ability to reabsorb these metabolites. Interestingly, we observed that SLC22A24 possesses a PDZ-consensus motif at its COOH-terminal end, which is also present in several known transporters localized to the apical membrane of the renal proximal tubule [73, 74] (**S11 Table**). The presence of this motif supports the direction of the observed associations and the idea that the transporter functions in the reabsorption of steroid conjugates in the kidney. To our knowledge, there have been no previously reported renal reabsorptive SLC transporters for steroid glucuronide conjugates, which have normally been found to be actively secreted in the kidney[75]. Future studies are required to confirm the membrane localization of the transporter in human kidney and to understand its physiological roles.

The highly significant association between the SLC22A24 nonsense variant and androsterone glucuronide levels led us to examine associations between the variant and diseases that are known to be associated with this metabolite. Through a phenome-wide scan of multiple publicly available sources, we observed associations between SLC22A24 and conotruncal heart defect (p=1×10^−6^)[76], lipid levels, and cardiovascular disease (p=1×10^−5^ to 9×10^−5^, data source Phenotype-Genotype Integrator, https://www.ncbi.nlm.nih.gov/gap/phegeni). In addition, we used phenome-wide association study (PheWAS) approach to analyze the many phenotypes in UKBiobank (http://geneatlas.roslin.ed.ac.uk/phewas/) with SLC22A24 SNPs that are associated with steroid and steroid conjugate levels, rs11231341 (p.Tyr501Ter) and rs78176967 (**S12 Table**). Measurement of hematological (e.g. immature reticulocyte fraction, neutrophil percentage) are among the top traits that are significantly associated with the two SNPs and these have been associated with steroid hormones[77–80] (**S12 Table**). Further, studies have shown that levels of androsterone glucuronide are elevated in women with acne and/or hirsutism[81–86]. Through a detailed search of associations with acne and/or hirsutism, we noted that the top SNPs in the SLC22A24 locus associated with acne (p=0.001) in the UKBiobank[87] are among the top SNPs significantly associated with steroid and steroid glucuronide levels (**S13 Table**). Overall, the identified phenotypic data consistent with the idea that dysregulation of steroid hormone metabolism contributes to the presentation of acne, blood traits, cardiovascular disease in humans (**S12 Table** and **S13 Table**).

In conclusion, these studies represent the first functional characterization of SLC22A24, a transporter in the UST (Unknown Substrate Transporter) region of the genome. The transporter has a unique substrate specificity and our studies suggest that it functions in the reabsorption of steroid glucuronide conjugates in the kidney (Fig 6C). This work represents an important step in the global effort to functionally characterize the remaining members of the SLC22 family.

## Materials and Methods

### Sources used to determine associations between genetic variations in SLC22A24 and human traits and diseases

Various public sources were used to obtain significant associations of SNPs within SLC22A24 gene +/− 50,000 kb (chr11:62553988-62743269 (hg18); chr11: 62797412-62986693 (hg19); chr11:63029940-63219221 (hg38)) with clinical variables related to metabolites levels. These sources included the GWAS Catalog, Phenotype-Genotype Integrator, GRASPS, GWAS Metabolic Server (http://metabolomics.gwas.eu) and others. The results from the search in public sources are shown in **S1 Table**.

### Further association analyses of steroid metabolite levels in the Atherosclerosis Risk in Communities Study (ARIC)

An application was approved by the ARIC Publication Committee to determine the associations of *SLC22A24* variants with steroid and bile acid metabolites in ARIC cohort (Manuscript proposal #3167). Below we described the ARIC study population used in this association analyses, metabolite measurement and association analyses of selected SNPs and metabolites relevant to this study. *Study population*: The ARIC is a prospective epidemiologic study conducted in U.S., designed to investigate the etiology and natural history of atherosclerosis, the etiology of clinical atherosclerotic diseases, and variation in cardiovascular risk factors, medical care and disease by race, gender, location, and date[88]. Using this cohort, investigators have published several papers related to associations between serum metabolome and diseases or traits among the subjects in the ARIC study[89]. In general, each visit included interviews and a physical examination [88]. In 2014, Metabolomic profiles were measured in baseline serum from 599 African- and 1,553 European-Americans. Participants were excluded if they did not give consent for use of DNA information. *Assessment of Metabolomic Profiles*: Methods for the assessment of metabolomics profiles, and whole exome sequencing were described previously [90–92]. In brief, metabolites were completed by Metabolon Inc. (Durham, USA) using an untargeted, gas chromatography-mass spectrometry and liquid chromatography-mass spectrometry (GC-MS and LC-MS)-based metabolomic quantification protocol. The annotated exome was captured by NimbleGen’s VCRome2.1 (Roche NimbleGen), and the captured exons were sequenced using Illumina HiSeq 2000. A total of 573 African Americans and 1375 Caucasians were available for the association analyses. The primary analysis was to determine the association of rs11231341 (SLC22A24 p.Tyr501Ter) with androsterone glucuronide (X-22379) and etiocholanolone glucuronide.

### Association of steroid glucuronide in Middle Eastern population

Yousri NA et al. have published the whole exome sequencing to identify genetic polymorphisms associated with metabolite levels in Middle Eastern population[9]. Using the data computed from Yousri NA et al.[9], the beta and p-value for the association of SLC22A24 p.Tyr501Ter (rs11231341) with androsterone glucuronide (X-22379) and etiocholanolone glucuronide are shown in Table 1.

### Clustering of human SLC22A24 and other species orthologs in the SLC22 family

Three phylogenetic trees were constructed. The first consists of the UniProt amino acid sequences for the human SLC22 family members with known substrates, human SLC22 members in the gene cluster between SLC22A8 and SLC22A9 of the human chromosome 11q12.3 (hg19 assembly), and all identified mouse and rat SLC22 members (Fig 2A). The second and third tree were built focusing on anion transporters. The second tree (Fig 4A) includes the human SLC22A24, known SLC22A24 orthologs in other species identified in the UniProt, known human anion transporters in SLC22 family (SLC22A6, A7, A8, A9, A11 and A12), genes in the human chromosome 11q12.3 (SLC22A10, A25), mouse anion transporters (Slc22a6, a7, a8, a9/a19, a9/a21, a12), mouse Slc22 genes in the clusters between Slc22a8 and Slc22a19 of the mouse chromosome 19qA (Slc22a26, a27, a28, a29, a30), rat anion transporters (Slc22a6, a7, a8, a9/a24, a12) and Slc22 genes in the clusters between Slc22a8 and Slc22a19 of the rat chromosome chr1q43 (Ust4r, Ust5r, Slc22a25). The third tree (**S4 Fig**) includes human SLC22A24 and the reference proteomes of all chordates (https://www.uniprot.org/help/reference_proteome), which have sequence similarity to human SLC22A24 with e-value of < 1×10^−200^. For each tree, a multiple sequence alignment was created using MUSCLE[93], with default parameters found on the CIPRES Science Gateway[94]. Phylogenetic trees were built using a RAxML[95] installation “RAxML-HPC2 on XSEDE” found on the CIPRES Science Gateway. The tree was calculated on protein sequences using the protein GAMMA model with the JTT substitution matrix and rapid bootstrapping with 100 replicas. A resulting extended majority rule consensus tree was visualized using Dendroscope 3[96]. See **S1 Appendix** and **S2 Appendix** for the amino sequences used to create the phylogenetic tree in Fig 2A, Fig 4A and **S4 Fig**.

### Comparative structure modelling

Due to the lack of available crystal structures, human OCT1 (SLC22A1), OCTN1 (SLC22A4), OAT1 (SLC22A6), and SLC22A24 were modeled using the crystal structure of the maltose-bound human GLUT3 (SLC2A3) in the outward-open conformation at 2.6 Å as a template (PDB ID 4ZWC)[97]. The template was selected on the basis of the shared MFS fold assignment [10], structure quality, and sequence similarity to the targets of interest (40%, 38%, 39% sequence similarity to SLC22A1, SLC22A6, SLC22A24, respectively). The sequence alignment was obtained by a manual refinement of gaps in the output from the PROMALS3D server (Figure SA1) [98]. The maltose molecule from the crystal structure was treated as the BLK residue type in the alignment and was copied from the template structure into a model as a rigid body. One hundred models were generated for each protein using the “automodel” class of MODELLER 9.16 (https://salilab.org/modeller/). The models had acceptable normalized discrete optimized protein energy scores (zDOPE)[99, 100]. The top scoring models were selected for electrostatic potential analysis with the APBS and PDB2PQR plugins[101] within UCSF Chimera using the AMBER force field with default settings[102]. Map values at surface points were exported using a custom Python script for subsequent orthosteric pocket analysis. Pocket electrostatic potential surface distributions were obtained by selecting all surface points within 7 Å (an average distance to the opening of the binding site as defined by POVME [103]) of any maltose HET atom.

### Establishment of transient and stable cell lines

Human embryonic kidney cell lines (HEK293) containing a Flp-In™ expression vector (HEK293 Flp-In) were used to create either transient or stable cell lines expressing SLC22A24 cDNA or vector only DNA. SLC22A24 (NM_173586 and NM_001136506) tagged open reading frame (ORF) clone and vector control (pCMV6-Entry) were purchased from OriGene. These vectors were either transiently transfected or stably transfected into Flp-In™ 293 cells using Lipofectamine LTX (Thermo Fisher Scientific). 500 ng of DNA and 1 µL of Lipofectamine LTX or 10 µg of DNA and 44 µL of Lipofectamine LTX were used for transfection into each well of the 24-well plate (seeding density 2.0×10^5^ cells/well) or 100 mm tissue culture plate (seeding density 4×10^6^ cells/well) respectively. More detailed methods to create stably or transiently transfected cells have been described from our groups (see[26, 38, 104]). For transient transfection, after 36 to 48 hours, cells were use for transporter studies (see section called Transporter uptake studies) or for protein quantifications (see section called Analysis of SLC22A24 protein levels in transiently expressing HEK293 cells.). For stable cell lines, after 48 hours, cells were transferred to a new 100 tissue culture plate and were treated with 500 µg/mL Geneticin^®^. Fresh media containing 500 µg/mL Geneticin^®^ was replaced each day.

### Cloning and expression profiling of SLC22A24 in liver and kidney

For the cloning, pooled total RNA of human kidney and liver were purchased from Clontech. Total RNA (2 µg) from each sample was reverse transcribed into cDNA using SuperScript VILO cDNA Synthesis kit (Thermo Fisher Scientific) according to the manufacturer’s protocol. For the expression profiling of SLC22A24, the cDNA from pooled human tissues was purchased from Takara Bio Inc. In particular the cDNA from two tissue panels was purchased, Human MTC^TM^ Panel I and II. **S9 Table** shows the primers that were used in the PCR to clone the transcripts, NM_173586 and NM_001136506. PCR products were cloned into pcDNA5FRT and sequenced (MCLAB, South San Francisco) to confirm the transcript. In addition, the primers that were used in the PCR to detect the presence of GAPDH (housekeeping gene) and the three SLC22A24 isoforms are shown in **S9 Table**. For expression profiling study, 10 ng cDNA of various tissues (Takara Human MTC panel I & II) were used for each reaction and amplified by KOD Xtreme Hot Start DNA polymerase kit (Takara). The PCR cycling conditions used are: (i) activation at 94°C for 2 min, (ii) denature at 98°C for 10 sec, (iii) annealing at 57.5°C for 30 sec and (iv) extension at 68°C for 1 min.

### RNA isolation and quantitative RT-PCR

HEK293 Flp-In cells were grown in 24-well poly-D-lysine coated plates (seeding density of 1.5-1.8 ×10^5^/well) until 75-80% confluency. Upon reaching confluency, cells were transiently transfected with vector only or vector containing different SLC22A24 isoforms (in pCMV6-Entry expression vector). The different isoforms were: SLC22A24 with 552 amino acids (NM_001136506.2, ENST00000612278.4), SLC22A24-551 with 551 amino acids (NM_001136506.2, ENST00000417740.5) and SLC22A24-322 with 322 amino acids (NM_173586.2) and SLC22A24-501Ter (c.1503T>G) (in both pCMV6-Entry and pcDNA5FRT vector backbone). 500 ng of the plasmid DNA, 1 µL of Lipofectamine LTX (Thermo Fisher Scientific) and 100 µL of the Opti-MEM I reduced serum media (Thermo Fisher Scientific) were used in the transfection mixture. 40 hours after transfection, media was removed and 350 µL of RNA Lysis buffer was added to each well. Total RNA was isolated using the Qiagen RNeasy kit (Qiagen). cDNA was synthesized using SuperScript VILO cDNA Synthesis Kit (Thermo Fisher Scientific). Quantitative RT-PCR (qRT-PCR) was performed using Taqman reagents and specific primer and probe sets for human SLC22A24 (Assay ID: Hs00543210_m) and GAPDH (Assay ID: Hs99999905_m1) (Applied Biosystems, Foster City, CA). Reactions were performed in a 96-well plate with a 10 µL reaction volume using an ABI 7900HT Fast Real-time PCR system (Applied Biosystems) using the default instrument settings. Expression levels were determined from three independent biological samples using the Ct method after normalization to endogenous levels of GAPDH[51, 105]. The results are expressed as a fold increase of the SLC22A24 mRNA level over the cell lines expressing the vector control.

### Synthesis of SLC22A24 ortholog cDNA

Sequences of SLC22A24 orthologs from mouse lemur (*Microcebus murinus*, Transcript ID ENSMICT00000042921.1), rabbit (*Oryctolagus cuniculus,* XM_002720992.3, Transcript ID ENSOCUT00000010486.2), and horse (*Equus caballus*, Transcript ID ENSECAT00000014835.1) were obtained from ensembl.org (release 95)[42] and the National Center for Biotechnology Information (NCBI). The cDNA of the gene was synthesized by GeneScript and inserted into the BamHI and XhoI restriction sites on the expression vector pcDNA5FRT (Invitrogen). The cDNA of the rat Slc22a9/Slc22a24 (NM_173302.1) was purchased from OriGene (Catalog number: RR202601). The synthesized SLC22A24 ortholog cDNA clones were used to make GFP fusion constructs to determine their subcellular localization. The GFP coding sequence was ligated to the 3’ end of the SLC22A24 ortholog cDNA in the expression vector pcDNA5/FRT using In-Fusion HD cloning kit (ClonTech #639642). The human SLC22A24 GFP fusion construct was purchased from OriGene (Catalog number: RG227944). The primers used to clone the GFP fusion constructs for the other non-human species are shown in **S9 Table**. The sequence of the constructs was confirmed by sequencing (MCLAB, South San Francisco).

### Transporter uptake studies

HEK293 Flp-In cells stably or transiently transfected with SLC22A24 were seeded at a density of 180,000 to 200,000 cells/0.5 mL in poly-D-lysine 24-well plates approximately 16 to 24 hours prior to uptake studies. For uptake studies in transiently expressing SLC22A24, methods described in previous section were performed before this study. Before the uptake studies, the culture medium (Dulbecco’s modified Eagle’s medium, DMEM H-21, supplemented with 10% fetal bovine serum) was removed and cells were incubated in 1.0 mL HBSS for 10-20 min at 37 °C. For the screening of radiolabeled compounds as substrate of SLC22A24, trace amount of the radiolabeled compounds (^3^H or ^14^C) were diluted in the HBSS (1:2000 or 1:3000) for uptake experiments. The details of the compound concentrations and uptake time are described in the results or figure legends. Uptake reactions were terminated by washing cells twice with 1 mL HBSS buffer and the cells were incubated in 700-800 µL of lysis buffer (0.1 N NaOH, 0.1% v/v SDS) and then 650-750 µL of cell lysate was transferred to scintillation fluid for scintillation counting. Experimental condition used in this study was similar to transporter assays published from our group[105, 106]. For pH dependence experiments, HBSS buffer was adjusted to different pHs (5.5, 7.4 and 8.5) by adding hydrochloric acid or sodium hydroxide[38]. For chloride dependence study, two different uptake buffer were used: (1) chloride free buffer (125 mM sodium gluconate, 4.8 mM potassium gluconate, 1.2 mM magnesium sulfate, 1.3 mM calcium gluconate, and 5 mM HEPES; adjusted with sodium hydroxide to pH 7.4); or (2) sodium buffer (125 mM sodium chloride, 4.8 mM potassium chloride, 1.3 mM calcium chloride, 1.3 mM magnesium sulfate, and 5 mM HEPES, adjusted with sodium hydroxide to pH 7.4)[105, 107]. For *trans*-stimulation studies, experimental conditions described from our published methods were used[38]. In brief, the SLC22A24 or EV stable cell lines were pre-incubated with either buffer or 2 mM succinic acid, 2 mM α-keto-glutaric acid or 2 mM glutaric acid for 2 hours. Then, the cells were washed twice with HBSS before initiating uptake of the anions (estrone sulfate, estradiol glucuronide, androstenediol glucuronide or taurocholic acid).

### Kinetic studies of steroid conjugates and bile acid transport by SLC22A24

We characterized the kinetics of the three steroid conjugates and two bile acids by SLC22A24, discovered from the initial screening. We selected SLC22A8 for comparison to SLC22A24 due to the significant accumulation of steroid conjugates by SLC22A8. The kinetic studies were performed following methods developed in our group[38]. In brief, these studies were performed using stable cell lines expressing SLC22A24 (NM_001136506) or SLC22A8[108]. For the kinetic studies of bile acid, only SLC22A24 expressing cells were used, as bile acids are poor substrates of human SLC22A8[109]. For each substrate, we varied the concentrations of the unlabeled compounds. The uptake rate was evaluated at 1 minute, as this fell within the linear range in the uptake versus time plot for each substrates. Each point represents the mean ± SD uptake in the transporter-transfected cells minus that in empty vector cells. The plots show the result for a representative experiment of n = 2.

### Inhibition of SLC22A24-mediated uptake by various anions and drugs

HEK293 Flp-In cells stably transfected with SLC22A24 (NM_001136506) were seeded at a density of 200,000 to 220,000 cells/0.5 mL in poly-D-lysine 24-well plates approximately 16 to 24 hours prior to the experiments. On the day of the experiments, cells were washed with 1 mL of Hank’s buffered salt solution (HBSS) per well and then preincubated for 10-20 min in the 1 mL of HBSS buffer. To assess inhibition, the cells were incubated with uptake buffer (1 µM unlabeled taurocholic acid and trace amount of [^3^H]-taurocholic acid in HBSS) containing either 100 µM estrone sulfate as positive control, 1% DMSO as negative control or 100 µM of the selected test compounds. Each of the 24-well plates contained negative and positive controls. Uptake of [^3^H]-taurocholic acid was stopped after 10 min by washing twice with ice-cold HBSS. The cells were lysed in 900 µL of lysis buffer (0.1 N NaOH and 0.1% SDS in double distilled water) per well while shaking for up to 90 min. 820 µL of the lysate was then added to 2.5 mL of EcoLite scintillation fluid (MP Bio), and the radioactivity was determined on a LS6500 Scintillation Counter (Beckman Coulter). Values were corrected for protein concentration as determined with a BCA assay kit (Thermo Scientific, Rockford, IL). All values were determined in triplicate, and final values are expressed as % uptake relative to negative control (1% DMSO). Methods used in the inhibition studies were from previous studies in our laboratory [110, 111]. The test compounds were tested at least one time and some compounds were selected to test two or three times to check for consistent results.

### Analysis of SLC22A24 protein levels in transiently expressing HEK293 cells

A 100 mm tissue culture plate of HEK293 Flp-In cells was transiently transfected with the following expression vectors: vector only (pCMV6-Entry), SLC22A24 (NM_001136506), SLC22A24-322 (NM_173586), SLC22A24-501Ter (created using site directed mutagenesis). All proteins contained a C-terminal Myc-DDK tag. For evaluation of the SLC22A24 protein levels, cell lysates were collected 48 hours after transfection. Plasma membranes were separated from intracellular membrane using the Plasma Membrane Protein Extraction Kit (Abcam ab65400)[33]. Transfected cells were harvested in 2 mL of homogenization buffer. Cells were homogenized using the syringe-based homogenization method. A needle (27 gauge) was attached to a 1 mL syringe. The cells lysates were passed through the syringe 5 times. A higher gauge needle, 30 gauge, was then used and the suspension was passed through for 5 additional times. The plasma membrane fraction was resuspended in 50 µL lysis buffer. Cell samples were deglycosylated using PNGase F (NEB P0708L) per the manufacturer’s protocol.

### Plasma membrane localization studies

The resulting GFP fusion constructs were transiently transfected in human embryonic kidney 293 (HEK293) cells. The cells were maintained in Dulbecco’s modified Eagle’s Medium (DMEM) (Fisher) supplemented with 10% fetal bovine serum (FBS) (Invitrogen) and 10 mM HEPES (Fisher). Dr. R. Irannejad’s laboratory has established standard methods for transfecting HEK293 cells for membrane localization studies[112–114]. HEK293 cells were grown to ∼70% confluence in poly-L-lysine coated glass coverslips in 12-well plates. The cells were transiently transfected with 1 µg plasmid DNA with 2 µL of Lipofectamine 2000 (Thermo Fisher Scientific). 24-hours after transient transfection, cells were fixed in 4% paraformaldehyde. Coverslips were mounted using ProLong Gold Antifade Mountant (Thermo Fisher Scientific) on glass microscope slides and visualized using Yokogawa CSU-22 Spining Disk Confocal microscope (Nikon Instruments, Inc).

### Metabolomic study

Samples of cell pellets from HEK293 Flp-In were prepared and sent to Metabolon Inc. (North Carolina, USA) for metabolites measurement using Metabolon^®^ platform. Cell pellet samples were prepared from five 100 mm culture dishes each of HEK293 Flp-In stably expressing vector only (pCMV6-Entry) or cells transiently expressing SLC22A24 cDNA (NM_001136506). 24 hours after transient transfection, media were removed and DMEM-H21, phenol red free, media (Invitrogen) containing 20% Fetal Bovine Serum, U.S.D.A. Approved Origin (Product Number 89510-186, Lot Number 356B16) were added to each dish. 48 hours after transient transfection, the cells were washed twice with cold PBS, and cells were scraped and transferred to a 15 mL falcon tubes. Tubes were centrifuged at 3000 rpm and supernatant were removed and stored the cell pellet in −80 °C freezer until shipping to Metabolon Inc. for metabolomics measurement (see[26] to obtain methods for metabolomic study).

### Metabolon data analysis

Raw metabolomics data were processed by Metabolon[115]. Individual metabolites were quantified and the values normalized. A two-tailed t-test was performed between EV and SLC22A24 and between EV and SLC22A1. False discovery rate (FDR) was used to correct p-values. We eliminated potential false positive metabolites using the following criteria. If the false discovery rate, FDR < 0.05 between SLC22A24 and EV cells but not significant (student t-test p-value 0.05) between SLC22A1 and EV cells, the metabolites were considered false positives. The results of the analysis are included in **S3 Table**.

### Site directed mutagenesis to create SLC22A24 p.Tyr501Ter

We used the Gibson Assembly protocol from New England BioLabs (catalog no. E5510) to create the SLC22A24 p.Tyr501Ter construct by amplifying unique segments from the SLC22A24 wild type and pCMV6-Entry vectors. The following primers were used to amplify the segment containing SLC22A24 residues 1-500 from the SLC22A24 wild type plasmid: forward primer GCCGCCGCGATCGCCATGGGCTTTGATGTGCTCCT, reverse primer CGGCCGCGTACGCGTGGAAATCCAGGGTAGGTGGG. The following primers were used to amplify the segment containing the linker, tag, stop codon, and the rest of the vector backbone from the pCMV6-Entry vector: forward primer CTACCCTGGATTTCCACGCGTACGCGGCCGCTCGA, reverse primer CACATCAAAGCCCATGGCGATCGCGGCGGCAGATC. Each segment contains a 15 bp overhang necessary for the Gibson Assembly workflow. The amplified segments were then assembled following the manufacturer’s protocol (https://www.neb.com/protocols/2012/12/11/gibson-assembly-protocol-e5510).

### Kidney tissue procurement

Frozen human postmortem frozen renal cortical tissues from donors aged 3 to 29 years old were obtained from the NIH NeuroBioBank at the University of Maryland, Baltimore, MD[116, 117]. These tissues were selected from patients, who are self-reported African American descent. Tissues were procured at the time of autopsy within 48 hours after death and were kept frozen at −80°C for later research. Tissues were selected for having no renal abnormalities in pathology and primary diagnosis. Twenty kidney samples were selected for quantitative proteomic analysis (see section below).

### Genotype kidney tissues for SLC22A24 c.1503T>G

Genomic DNA from the 20 renal cortical tissues procured from the NIH NeuroBioBank were isolated using Wizard genomic DNA purification kit (Promega). The following forward (CCACAAGGGCAGAAAGTATG) and reverse primers (CAAATTATCAAAGCTGCGGGTTA) were used to PCR the genomic region of SLC22A24 and using Sanger sequencing primer (TAAGCCAGATATTGTTCACG) to determine the genotype for the stop-gained variant in SLC22A24, c.1503T>G (rs11231341).

### Quantitative proteomics

Absolute quantitative proteomics was performed in the 20 renal cortical tissues. SLC22A24 and SLC22A24-551 were quantified using LC-MS/MS with Selective Reaction Monitoring (SRM) approach by Omics Technologies, Inc. (http://www.proteomics.omicstech.com/proteomics/srm.html, previously known as MyOmicsDx, Inc., Towson, MD, USA). The following are the main steps for quantitative proteomic analysis for the three transcript isoforms of SLC22A24. The detailed material and method used for each steps are described below as well as in our previous submitted manuscript[117]. The material and methods described below were provided by Omics Technologies, Inc..

- Membrane protein extraction and protein preparation
- Chemicals and Reagents
- Preparation of the Chromatography Solutions
- SRM analysis
- Tuning of the Agilent 6495 Triple Quadrupole mass spectrometer
- SRM data analysis

#### • Membrane protein extraction and protein preparation

Frozen renal cortical tissues were processed to extract membrane proteins. Membrane protein samples were then processed by Omics Technologies, Inc. using “Filter Assisted Sample Preparation” (FASP) method[118]. Briefly, protein samples in 9M UREA were reduced with 5 mM TCEP at 37 °C for 45 min and reduced cysteine were blocked using 50 mM iodoacetmide (IAA) at 25 °C for 15 min. Protein samples were then cleaned using 10 kDa Amicon Filter (UFC501096, Millipore) three times using 9M urea and two times using MyPro-Buffer 1 (Omics Technologies, Inc.). Samples were then proteolyzed with trypsin (V5111, Promega) for 12 hrs at 37 °C. The peptide solution then was acidified by adding 1% trifluoroacetic acid (TFA) and was incubated at room temperature for 15 min. A Sep-Pak light C18 cartridge (Waters Corporation) was activated by loading 5 mL 100% (vol/vol) acetonitrile and was washed by 3.5 mL 0.1% TFA solution two times. Acidified digested peptide solution was centrifuged at 1,800 × g for 5 min, and the supernatant was loaded into the cartridge. To desalt the peptides bound to the cartridge, 1 mL, 3 mL, and 4 mL of 0.1% TFA were used sequentially. To elute the peptides from the cartridge, 2 mL of 40% (vol/vol) acetonitrile with 0.1% TFA was used. The eluted peptides were lyophilized overnight and reconstituted in 37 μL MyPro-Buffer 3 (Omics Technologies, Inc.).

#### • Chemicals and Reagents

TCEP (TCEP (tris-(2-carboxyethyl) phosphine) and MMTS (methyl methanethiosuphonate) were purchased from Thermo Scientific (Waltham, Massachusetts). LysC and Trypsin proteases were purchased from Promega (Fitchburg, Wisconsin). C18 Cartridges for sample preparation, and chromatography columns for basic reverse phase liquid chromatographic (bRPLC) and online HPLC of Triple Quadrupole mass spectrometer were purchased from Waters (Milford, Massachusetts). Acetonitrile was purchased from JT Baker, and formic acid was obtained from EMD Millipore (Billerica, MA, USA). MyPro-Buffer 1, MyPro-Buffer 2 and MyPro-Buffer 3 were utilized by Omics Technologies, Inc. All other reagents were purchased from Sigma-Aldrich (St. Louis, Missouri) unless otherwise indicated.

#### • Preparation of the Chromatography Solutions

bRPLC Solvent A contained 10mM triethylammonium bicarbonate buffer (TEABC); bRPLC Solvent B contained 10mM TEABC, 90% (vol/vol) Acetonitrile. SRM Mass Spectrometry Solvent A was comprised of water with 0.1% (vol/vol) Formic Acid; SRM Solvent B was Acetonitrile with 0.1% (vol/vol) Formic Acid.

#### • SRM analysis

Peptide samples reconstituted in 37 µl MyPro-Buffer 3 (MyOmicsDx, Inc) were spiked with MyPro-SRM Internal Control Mixture (MyOmicsDx, Inc) composed of a pool of 1 femto mole heavy isotope labeled peptides covering a large hydrophobicity window and a large M/z range (M/z 200 ∼ 1300), and were subject to SRM analysis. Peptide samples were eluted through an online Agilent 1290 HPLC system into the Jet Stream ESI source of an Agilent 6495 Triple Quadrupole Mass spectrometer.

#### • Tuning of the Agilent 6495 Triple Quadrupole mass spectrometer

After every preventative maintenance, Agilent 6495 Triple Quadrupole mass spectrometer was tuned using manufacturer’s tuning mixture followed by MyPro-SRM Tuning Booster (MyOmicsDx SRM tuning mixture). Before and after every batch of SRM analysis, to ensure the stable and consistent performance of the mass spectrometer throughout the entire study, MyPro-SRM Performance Standard (MyOmicsDx), a mixture of standard peptides across a wide range of mass (M/z 100-1400) and a broad range of hydrophobicity were analyzed.

#### • SRM Data analysis

The abundance of a target peptide was represented by the total area under the curve (AUC) of all its transitions normalized to the total AUC of all transitions from the most nearby (sharing a similar hydrophobicity) heavy isotope-labeled peptide from MyPro-SRM Internal Control Mixture (MyOmicsDx, Inc) spiked in before the SRM analysis. Absolute quantification of each protein is performed through applying AQUA™ Peptides purchased from Sigma-Aldrich. A summary of all quantification results were shown in **S8 Table**.

## Acronyms

SLC: solute carrier
ABC: ATP-binding cassette
GWAS: genome-wide association studies
SNP: single nucleotide polymorphism
UST: unknown substrate transporter
ES: estrone sulfate
DHEAS: dehydroepiandrosterone sulfate
TA: taurocholic acid
GA: glycocholic acid
EG: estradiol glucuronide
AG: androsterone glucuronide
SA: succinic acid
6-CF: 6-carboxyfluorescein
DBF: 4’,5’-dibromofluorescein
DCF: 2’,7’-dichlorofluorescein
EV: empty vector

## Acknowledgements

This work was supported by the National Institutes of Health [GM117163 and DK108722 to K.M.G. and S.W.Y.]. The Atherosclerosis Risk in Communities study has been funded in whole or in part with Federal funds from the National Heart, Lung, and Blood Institute, National Institutes of Health, Department of Health and Human Services (contract numbers HHSN268201700001I, HHSN268201700002I, HHSN268201700003I, HHSN268201700004I and HHSN268201700005I). The authors thank the staff and participants of the ARIC study for their important contributions. Funding support for “Building on GWAS for NHLBI-diseases: the U.S. CHARGE consortium” was provided by the NIH through the American Recovery and Reinvestment Act of 2009 (ARRA) (5RC2HL102419). Metabolomics measurements were sponsored by the National Human Genome Research Institute (3U01HG004402-02S1). Sequencing was carried out at the Baylor College of Medicine Human Genome Sequencing Center and supported by the National Human Genome Research Institute grants U54 HG003273 and UM1 HG008898. K.S. and N.A.Y. were supported by the Biomedical Research Program funds at Weill Cornell Medical College in Qatar, a program funded by the Qatar Foundation. The authors thank the National Institute of Health (NIH) Neurobiobank at the University of Maryland, Baltimore, M.D. for providing the frozen human kidney tissue. The authors also thank the Omics Technologies Inc. for their services to quantify the SLC22A24 protein levels in the renal cortical tissues and Dr. Tao Long, for personal communications regarding the interpretation of the GWAS metabolomic data[8].

## S1 to S9 Figure Legend

**S1 Figure.** A common splice donor variant, rs1939749, found in 3’-end of exon 1. **A.** A splice donor, rs1939749, C>T (+ DNA strand), is strongly associated with lower expression levels of SLC22A24 in kidney tubular cells observed in two independent kidney eQTL databases, http://susztaklab.com/eqtl and http://nephqtl.org/ and p-values 3.7×10^−23^ (beta = 1.2, N=121) and 1.9×10^−13^ (beta = 0.79, N=166) respectively. **B.** Zoom in of the SLC22A24 gene region (hg19) flanking exon 1 and exon 2 with padding of 25 bases. This splice donor, rs1939749, C>T (+ DNA strand), creates a premature stop.

**S2 Figure.** Kinetic of uptake of androstenediol glucuronide for SLC22A24. The uptake rate was evaluated at 2 minutes. Each point represents the mean ± S.D. uptake in the SLC22A24-stably transfected cells minus that in empty vector cells. Figure shows a representative plot from one experiment. The kinetic parameters of androstanediol-3-glucuronide could not be determined accurately due to a solubility limit of 300 µM. Using the plateau at 300 µM as an approximation, the affinity of androstanediol glucuronide was calculated as 741 ± 253 μM. The data were fit to a Michaelis−Menten equation.

**S3 Figure.** Effect of sodium and pH on SLC22A24-mediated uptake of anion. **A.** ^3^H-Estrone sulfate (with 1 µM unlabeled estrone sulfate) was incubated in the Na+ buffer (125 mM sodium chloride, 4.8 mM potassium chloride, 1.3 mM calcium chloride, 1.3 mM magnesium sulfate, and 5 mM HEPES, adjusted with sodium hydroxide to pH 7.4) or Na+ free buffer (125 mM choline chloride, 4.8 mM potassium chloride, 1.3 mM calcium chloride, 1.3 mM magnesium sulfate, and 5 mM HEPES, adjusted with TRIS-HCl to pH 7.4) for 10 min. Data shown are from the mean ± S.D. for representative experiment of n=2. **B.** ^3^H-Esterone sulfate, ^3^H-estradiol glucuronide or ^3^H-taurocholic acid (with 1 µM unlabeled compound) was incubated in the HBSS buffer adjusted to pH 5.5, 7.4 and 8.5, and the cells were incubated for 10 min. Data shown are from the mean ± S.D. for representative experiment of n=2.

**S4 Figure.** Phylogenetic tree of indicating protein sequence relationships between human USTs and identified mammalian orthologs. The unique and stable entry identifier from UniProtKB (https://www.uniprot.org/) are shown in parentheses. Red arrow points to the human SLC22A24.

**S5 Figure.** Plasma membrane expressions of SLC22A24-GFP tagged from five different species. GFP fusion constructs were generated for the human, rabbit, horse, and mouse lemur SLC22A24 and the rat Slc22a9/Slc22a24. HEK293 cells were transient transfected with the GFP fusion constructs. The cells were visualized by confocal microscopy to determine whether the GFP fused protein is expressed on plasma membrane.

**S6 Figure.** Plasma membrane expression, transport uptake and predicted structure of human SLC22A24 or SLC22A24-322. **A.** GFP fusion constructs were generated for human SLC22A24 and SLC22A24-322. HEK293T cells were transient transfected with the GFP fusion constructs. The cells were visualized by confocal microscopy to determine whether the GFP fused protein is expressed on plasma membrane. **B.** The uptake experiments of various anions were performed for 10 minutes in triplicate wells using trace amount of the radiolabeled compound. Experiments were repeated once. **C.** A comparative structure model of the human SLC22A24 with 12 transmembrane helices (white and red). SLC22A24-322 is predicted to consist of transmembrane helices 1-6 (white), but missing transmembrane helices 7-12 (red).

**S7 Figure.** Schematic figure showing the amino acid differences between SLC22A24 and its isoform, SLC22A24-551. **A.** Zoom in of the SLC22A24 gene region (reverse direction) between exon 9 and exon 10 along with padding of 10 bases using UCSC browser. The allele “CT” from position 1 and position 2 resulted in formation of two different isoforms (splice variants) of SLC22A24. **B**. Zoom in of the SLC22A24 amino acids between exon 9 and exon 10. Arrows show the differences in amino acid positions 533 and 534 and hence the SLC22A24 and SLC22A24-551 protein.

**S8 Figure.** Protein and transcript levels of SLC22A24 quantified by western blotting and qRT-PCR using Taqman assay in HEK293-FlpIn cells transiently expressing empty vector, SLC22A24, SLC22A24-322 and SLC22A24-501Ter. Western blotting (**A**) showed that SLC22A24 non-sense variant, SLC22A24-501Ter and SLC22A24-322 do not expressed on the plasma membrane despite mRNA levels are detectable (**B**).

**S9 Figure.** The peptides of interest and their corresponding fragments to quantify the protein levels of SLC22A24 and SLC22A24-551. Peptide 1 corresponds to the SLC22A24 and SLC22A24-551 proteins; whereas peptide 2 and 3 correspond specifically to the SLC22A24 and SLC22A24-551 respectively.

## S1 to S13 Table Legend

**S1 Table**. Metabolites that are significantly associated with SNPs within SLC22A24 locus (chr11:62553988-62743269 (hg18); chr11: 62797412-62986693 (hg19); chr11:63029940-63219221 (hg38)). The results are sorted by p-value.

**S2 Table.** Association of SLC22A10 and SLC22A25 missense and nonsense variants with two steroid conjugates. These functional SNPs have weaker association with steroid glucuronide compared to SLC22A24 (501-Ter, rs11231341) (S1 Table).

**S3 Table**. Results from metabolomic study using Metabolon Inc. technology. Values for each sample are normalized by Bradford protein concentration. Each biochemical is then rescaled to set the median equal to 1. Missing values are imputed with the minimum. Values from columns N to R, columns S to W and columns X to AB are values from HEK293 Flp-In cells transient transfected with empty vector (EV), SLC22A1 (positive control) and SLC22A24. Columns AV to AE are calculated average from five samples transfected with EV, SLC22A1 or SLC22A24. Student t-test was used to determine the p-values between EV and SLC22A1 or EV and SLC22A24 (columns AF and AG). False discovery rate (FDR) was calculated, as shown in columns AI and AJ.

**S4 Table.** Experimental data of uptake in HEK293-FlpIn cells transfected by SLC22A24, empty vector or other transporters in the SLC22 family, which are used as positive control for the specific substrate tested. n.s = fold uptake between EV and SLC22A24 < 1.5-fold and t-test p-value >0.05.

**S5 Table. Kinetic parameters of steroid conjugates and bile acids uptake by SLC22A24.** Kinetic parameters of the estrone sulfate, estradiol glucuronide, androstanediol glucuronide and taurocholic acid by SLC22A8 are included here as comparison to SLC22A24. The values are average ± SD from two experiments.

**S6 Table.** List of metabolites or prescription drugs and their metabolites, which were screened for inhibition of SLC22A24-mediated 3H-taurocholic acid uptake (Figure 3g).

**S7 Table**. The cDNA from two tissue panels were purchased from Takara Bio Inc. Human MTC^TM^ Panel I and II contained sixteen different tissues.

**S8 Table.** SRM quantification of different isoforms of protein SLC22A24, using three different peptides in 20 renal cortical samples.

**S9 Table**. Primers used for PCR to detect SLC22A24 transcripts and to clone the SLC22A24 cDNA. PCR conditions are also provided in table below.

**S10 Table.** Transcript levels of 22 genes in SLC22 family in kidney samples available from two databases HPA (https://www.proteinatlas.org) and GTEx (https://www.gtexportal.org/).

**S11 Table**. List of renal transporters in the solute carrier (SLC) superfamily that are known to have apical membrane localization. It has been demonstrated that apical membrane localization transporters possess a PDZ-motif consensus (S/T-X-h, where X is any amino acid and h is hydrophobic amino acids (A, F, L, M, I, W, P and V)) at their COOH-terminal end. SLC22A24 has the PDZ-domain consensus sequence and support its role on apical membrane of the kidney.

**S12 Table.** Pheno-wide association results of two SNPs in SLC22A24 that are associated with steroid and steroid conjugate levels (rs11231341 and rs78176967) with human disease and traits in UKBiobank (http://geneatlas.roslin.ed.ac.uk/phewas/).

**S13 Table.** Association results of variants in SLC22A24 locus (chr11:62,799,410-62,986,182, hg19) with acne/acne vulgaris extracted from UKBiobank (http://biobank.ctsu.ox.ac.uk/crystal/field.cgi?id=20002 (N=457 cases)). The summary statistic for acne/acne vulgaris was obtained from Neale Lab (http://www.nealelab.is/uk-biobank/). Allele2 (column P) is the effect allele. A positive beta means that the allele2 (effect allele) is associated with increased risk of acne/acne vulgaris.

**S1 Appendix**. FASTA format representing amino acid sequences of solute carrier family 22 (human, mouse and rat). These sequences in FASTA format is used to create phylogenetic tree in Fig 2A.

**S2 Appendix.** FASTA format representing amino acid sequences of anion transporters of the solute carrier family 22 (human, primates, mouse, rat and other species). These sequences in FASTA format is used to create phylogenetic tree in Fig 4A and **S4 Fig**.

